# A coarse-grained resource allocation model of carbon and nitrogen metabolism in unicellular microbes

**DOI:** 10.1101/2023.04.04.535571

**Authors:** Istvan T. Kleijn, Samuel Marguerat, Vahid Shahrezaei

## Abstract

Coarse-grained resource allocation models (C-GRAMs) are simple mathematical models of whole cell physiology, where large components of the macromolecular composition are abstracted into single entities. The dynamics and steady-state behaviour of such models provides insights on optimal allocation of cellular resources and have explained experimentally observed cellular growth laws. Here, we formulate a minimal C-GRAM with nitrogen and carbon pathways converging on biomass production, with parameterizations accounting for respiro-fermentative and purely respiratory growth. The model describes the effects of the uptake of sugars, ammonium, and/or compound nutrients such as amino acids on the translational resource allocation towards proteome sectors that maximised the growth rate. It robustly recovers cellular growth laws including the Monod law and the ribosomal growth law. Furthermore, we show how the growth-maximizing balance between carbon uptake, recycling, and excretion depends on the nutrient environment. Lastly, we find a robust linear correlation between the ribosome fraction and the abundance of amino acid equivalents in the optimal cell, which supports the view that simple regulation of translational gene expression can enable cells to achieve an approximately optimal growth state.

## 1 Introduction

Unicellular organisms are remarkably efficient self-replicators as they are under selective pressure to grow fast or risk being outcompeted by rival colonies or species (Dekel and Alon 2005; Lynch et al. 2014). On the other hand, microbial cells are faced with internal constraints limiting their growth, because each metabolite, macromolecule, or unit of membrane area can only be used for one reaction at any given time. Fast-growing cells must therefore possess an ability to allocate these limited resources in varied environments (Koch 1988; López-Maury et al. 2008; Berkhout et al. 2013; Bruggeman et al. 2020). The interplay between gene expression and growth can be studied reproducibly in states of balanced growth, where cells are maintained in the same environment for many generations (Schaechter 2006).

Observed patterns of gene expression and the resulting rate of growth depend heavily on the growth environment, in particular on the presence of external stresses and, importantly, the nutrient makeup. A well-established feature of gene expression in multiple model organisms is the presence of linear correlations between broad classes of macromolecules and the cellular growth rate (Schaechter et al. 1958; Mitchison and Lark 1962; Waldron and Lacroute 1975; Bremer and Dennis 2008; Zavřel et al. 2019). The most pervasive positive correlation is between the abundance of translational resources (chiefly ribosomes) and the growth rate (Scott et al. 2010; You et al. 2013; Schmidt et al. 2016; Metzl-Raz et al. 2017; Kleijn et al. 2022). Classes of proteins that are negatively correlated with the growth rate are associated with stress, or induced by the specific cause of growth inhibition—stressors or reductions in the nutrient quality or quantity (Brauer et al. 2008; Hui et al. 2015; Schmidt et al. 2016; Kleijn et al. 2022).

The concept of proteome sectors has been the basis of several coarse-grained phenomenological and mechanistic models relating optimal resource allocation to protein abundance and cellular growth rates (Molenaar et al. 2009; Scott et al. 2014; Maitra and Dill 2015; Weiße et al. 2015; Maitra and Dill 2016; Pandey and Jain 2016; Liao et al. 2017; Bertaux et al. 2020; Hu et al. 2020). In coarse-grained resource allocation models (C-GRAMs), large sectors of the proteome are abstracted into a single protein, whose kinetics are explicitly described. Extended summaries of earlier models are provided in Supp. Text 5.1. The coarse-grained approach allows for direct interpretation of key model parameters, such that the model can be explored directly with minimal need for explicit parameter inference.

Earlier coarse-grained modelling efforts chiefly considered carbon modulations representative for the effect of the nutrient quality in general. In such models, metabolism was typically considered as a linear pathway from nutrient to protein production. This included the models proposed by Weiße et al. (2015) and by Bertaux et al. (2020). Contrasting with this earlier work, a strategy commonly employed in *Schizosaccharomyces pombe* to modulate the growth rate uses ammonium chloride and a variety of amino acids as nitrogen sources (Carlson et al. 1999; Kleijn et al. 2022). Using this strategy, we have recently reported differential expression for many enzymes involved in carbon metabolism across nitrogen sources, even though abundant glucose was provided in all conditions (Kleijn et al. 2022). Different amino acids have also been used as sole nitrogen sources in order to modulate growth while studying the proteome of *Saccharomyces cerevisiae* (Yu et al. 2021) and of three bacterial strains found in the *Arabidopsis* rhizosphere (Jacoby et al. 2020).

Here, we aim to better understand the effect of nitrogen-source modulation on resource allocation in a coarse-grained modelling context. Molenaar et al. (2009) constructed a minimal C-GRAM that incorporated nitrogen metabolism and described proteome allocation under optimal growth. However, they did not account for the uptake of complex nitrogen sources such as amino acids. More recently, Bertaux et al. (2020) took a similar approach and additionally incorporated metabolites in the calculation of the system size. We opted to extend the latter to include the uptake and metabolism of carbon-containing nitrogen sources. In this modelling paradigm, we studied how a growth-rate-maximising allocation towards proteome sectors varied between steady states that were imposed by choices of parameters representing different nutrient conditions.

## 2 Results

In this study, we aim to investigate optimal resource allocation behaviour under growth in defined media with varying nitrogen sources. This for example, will be relevant for growth media where different amino acids act as the sole source of nitrogen. Amino acids generally consist of an amino group and a ketoacid backbone, and we therefore focused on modelling nitrogen and carbon metabolism. We constructed a coarse-grained resource allocation model (C-GRAM) with pathways representing nitrogen and carbon uptake, protein biosynthesis, and the excretion and recycling of carbon from carbon-containing nitrogen sources. The model was formulated dynamically using ordinary differential equations (ODEs), their steady-state representing balanced growth. The steady-state growth rate was calculated from the solution to the ODEs as an emergent property.

Pathways in the model were represented by enzymes with simple kinetics, and the parameters in this representation were chosen to reflect nutrient conditions. For example, carbon limitation was modelled by reducing the catalytic rate of the carbon uptake enzyme. Furthermore, nitrogen sources were distinguished by the catalytic rates of the enzymes metabolising them as well as their stoichiometries, related to their elemental carbon-to-nitrogen ratio. For each nutrient condition (parameterization), we determined the resource allocation that maximised the growth rate. As expected, various nutrient limitations induced complex trade-offs between allocations to different enzyme fractions. The details regarding the model implementations, namely the full formulation of the ODEs, considerations with regards to parameter choices, and the approach used to optimise resource allocation are described in the Methods.

As described in the remainder of this manuscript, we explored the effect of different growth-rate modulations on the optimal allocation. We first considered the behaviour of two submodels that lacked the recycling and excretion of the ketoacid backbone. This enabled us to explore trade-offs in resource allocation in media with simple sources of carbon (such as glucose) and nitrogen (such as ammonium chloride). In the first submodel, only one type of metabolism was explored; in the second submodel, respiratory metabolism was introduced. Finally, the full model was used to describe growth on amino-acid nitrogen sources.

### 2.1 Modulation of the carbon uptake rate with one metabolic pathway

The first submodel we explored was one with parallel uptake of carbon and nitrogen, which were combined by a single amino-acid-producing enzyme. An illustration of the model is provided as Fig. 1A. This model was explored by varying the carbon uptake rate parameter k_C_.

**Figure 1:**
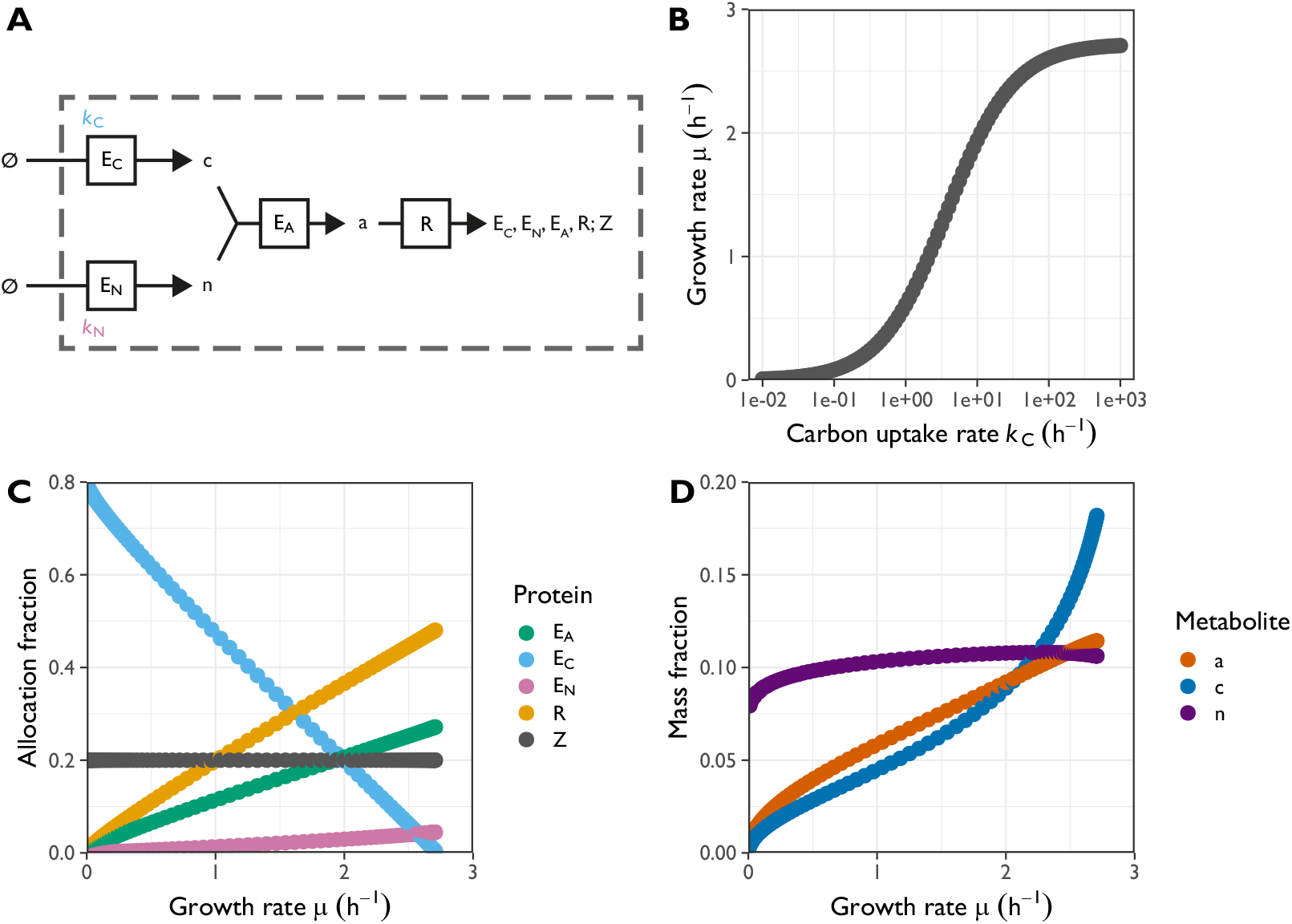
Analysis of the core metabolic model under modulation of the carbon uptake rate k_C_. The uptake rate parameter k_C_ of the transporter was modulated to yield different growth rates. The allocation fractions were chosen to maximise the growth rate for each chosen k_C_. **A.** Illustration of the model. **B.** Growth rate μ as a function of the carbon uptake rate k_C_. **C.** Optimal protein allocation fractions as a function of the growth rate μ. **D.** Steady-state mass fractions of metabolites as a function of the growth rate μ.

As expected, the dependency of the growth rate on the modulation parameter approximated a Monod curve (Fig. 1B), and the optimal protein allocation fractions were approximately linearly correlated with the growth rate (Fig. 1C). The total relative metabolite abundance in steady-state amounted to approximately 10%-30% of biomass (Fig 1D). In experimental cultures, metabolites comprise only a small fraction of the biomass, so in this respect the growth-optimised model was a poor approximation for conditions where this occurred. This may reflect the current parameterization of the model to be inaccurate.

Importantly, under modulation of the carbon uptake enzyme, the nitrogen assimilation enzyme E_N_ and the amino-acid biosynthesis enzyme E_A_ were part of the “R”-fraction: their allocation fractions were positively correlated with the growth rate (Fig. 1C). This behaviour was also implemented by Molenaar et al. (2009), but it is notably different from the model described by Weiße et al. (2015), who assumed that the transporter and metabolic enzyme were both regulated identically.

Proteins associated with translation are estimated to constitute approximately 45% of the total protein mass in the fastest-growing *E. coli* cultures (Scott et al. 2010; Hui et al. 2015; Schmidt et al. 2016), which reach maximal growth rates of around 2.0 h^-1^. A minimal partitioning of the proteome based on growth rate correlations suggested that around half of the proteome mass does not change with the growth rate (Scott et al. 2010, 2014). The earlier model by Weiße et al. (2015) reflected this by implementing protein expression regulation with a metabolic sector, a translational sector, and a constitutive sector. Its proteome comprised two sequential enzymes (corresponding to our E_C_ and E_A_), ribosomes (our R), and housekeeping proteins (our Z). In their parameterization, the latter took up around 70% of the total protein mass, and were negatively correlated with the growth rate.

In contrast, in the parameterization used to generate Fig. 1, the allocation towards housekeeping proteins was only 20% of the proteome. However, this para meterization was chosen such that the maximal ribosomal allocation agreed reasonably well with *E. coli* data from Scott et al. (2010). The discrepancy between the two models in their estimated housekeeping allocations is explained by the considerable cost of the non-modulated enzymes. At maximal growth, the aminoacid synthesis and nitrogen-uptake enzymes amount to around 25% and 5% of the proteome, respectively.

### 2.2 Modulation of carbon and nitrogen uptake rates with two parallel metabolic pathways

Next, we considered that a key determinant of unicellular growth is the metabolic state, i.e. whether fermentation occurs or whether biosynthesis occurs in a purely respiratory fashion. We wondered under what circumstances such changes in metabolism would be triggered and therefore implemented the model with two parallel amino-acid synthesis pathways. These parallel pathways differed only in their parameterizations, with the purely respiratory enzyme E_Ar_ requiring less carbon substrate, but a higher expression to sustain the same synthesis rate, than the respirofermentative enzyme E_Af_ (Methods section 4.3). The allocation fractions to both enzymes were included in the growth rate optimisation. We modulated the carbon and nitrogen transporter rate parameters in the model containing two parallel metabolic pathways with different parameterizations (see Fig. 2A).

**Figure 2:**
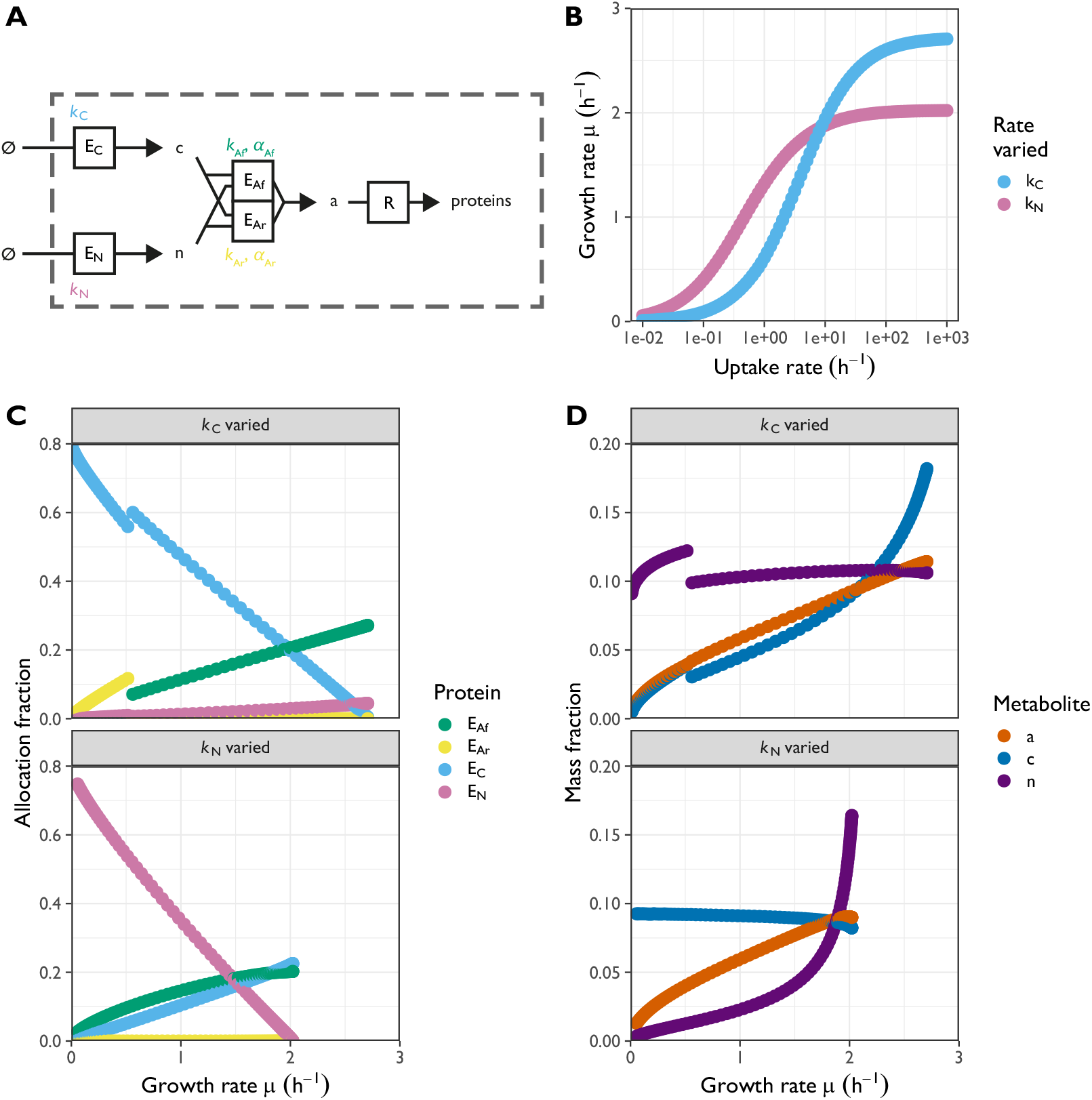
Analysis of the model with two parallel metabolic enzymes (respiro-fermentative enzyme E_Af_ and purely respiratory enzyme E_Ar_) under the separate modulation of the carbon and nitrogen transporter uptake rates k_C_ and k_N_, respectively. Allocation fractions were chosen to maximise the growth rate for each chosen kC (C. and D., top panels) and k_N_ (bottom panels). The uptake rates were varied separately between 1.0 × 10^-2^ h^-1^ and 1.0 × 10^3^ h^-1^. **A.** Illustration of the model. **B.** Growth rate μ as a function of the respective uptake rate. **C.** Optimal protein allocation fractions as a function of growth rate μ for the two parameter explorations. Ribosomes R and housekeeping proteins Z were omitted from the figure to improve clarity. **D.** Steady-state mass fractions of metabolites as a function of growth rate μ.

As before, the growth rate depended on the modulation parameters approximately following a Monod curve (Fig. 2B). Because the two pathways are parallel, the usage of one of the two was preferred over the other in each parameterization, such that one of E_Ar_ and E_Af_ was expressed while the other was not. As shown in Fig. 2C, the purely respiratory enzyme was induced by low values of the carbon uptake rate k_C_. We note a sharp and discontinuous transition from respiration to fermentation at a specific level of carbon uptake rate (Fig. 2C). This agrees with results of a model similar in scope to ours, where the optimal choice between a metabolically efficient or catabolically efficient pathway was shown to depend on the growth rate (Molenaar et al. 2009). In contrast to this behaviour under modulations of the carbon uptake, the respirofermentative enzyme was present in all conditions where the nitrogen uptake efficiency k_N_ was varied from its default.

Note that purely respiratory metabolism required a higher abundance of both raw metabolites, and therefore both c and n varied discontinuously with the growth rate near the transition point (μ ≈ 0.5 ^-1^h). However, the amino acid abundance a varied almost continuously with the growth rate even at the transition point. The relation between a and the ribosomal allocation will be explored in detail in a further section of this paper.

Under perturbations of the nitrogen transporter uptake rate k_N_ (Fig. 2C bottom panel), the allocation towards the metabolic enzyme varied nonlinearly with the growth rate. In addition to this, the amount of nitrogen metabolite n built up much more strongly with increasing growth rate via k_N_ than the equivalent carbon build-up under perturbations of k_C_ (Fig. 2D). Note that modulating each transporter’s efficiency (k_N_ and k_C_) repressed the abundance of its own substrate (n and *c* respectively). On the other hand, the modulation of one transporter rate barely affected homeostasis of the metabolite in the other pathway (c and n, respectively).

### 2.3 Recycling and excretion of ketoacids disturbed carbon metabolism

In the above, we restricted ourselves to the metabolism of freely usable nitrogen, which typically comes in the form of ammonium ions (NH_4_^+^). However, the cellular growth rate can also be perturbed considerably by using different nitrogen sources, particularly amino acids, as the sole nitrogen source instead of NH_4_^+^ (Carlson et al. 1999; Kleijn et al. 2022). These pathways typically de- or transaminate the amino acid, either by a single enzyme or as the net result of a longer pathway (Godard et al. 2007). Nitrogen is assimilated as free NH_4_^+^ (after deamination) or glutamate (after transamination). The leftover carbohydrate, usually a ketoacid, may be recycled into biomass or excreted.

To account for this process in the model, we implemented the nitrogen uptake pathway as an enzyme that produced both free nitrogen and a carbon-containing ketoacid metabolite. To represent different complex nitrogen sources such as amino acids, we perturbed the parameter γ_K_, which represents the relative mass of recycled ketoacid with respect to the total mass taken up by the nitrogen uptake enzyme. Additionally, we introduced two enzymes, E_Kre_ and E_Kex_ that, respectively, recycled the ketoacid into usable carbon precursor or excreted it from the cell. The model is illustrated in Fig. 3A.

**Figure 3:**
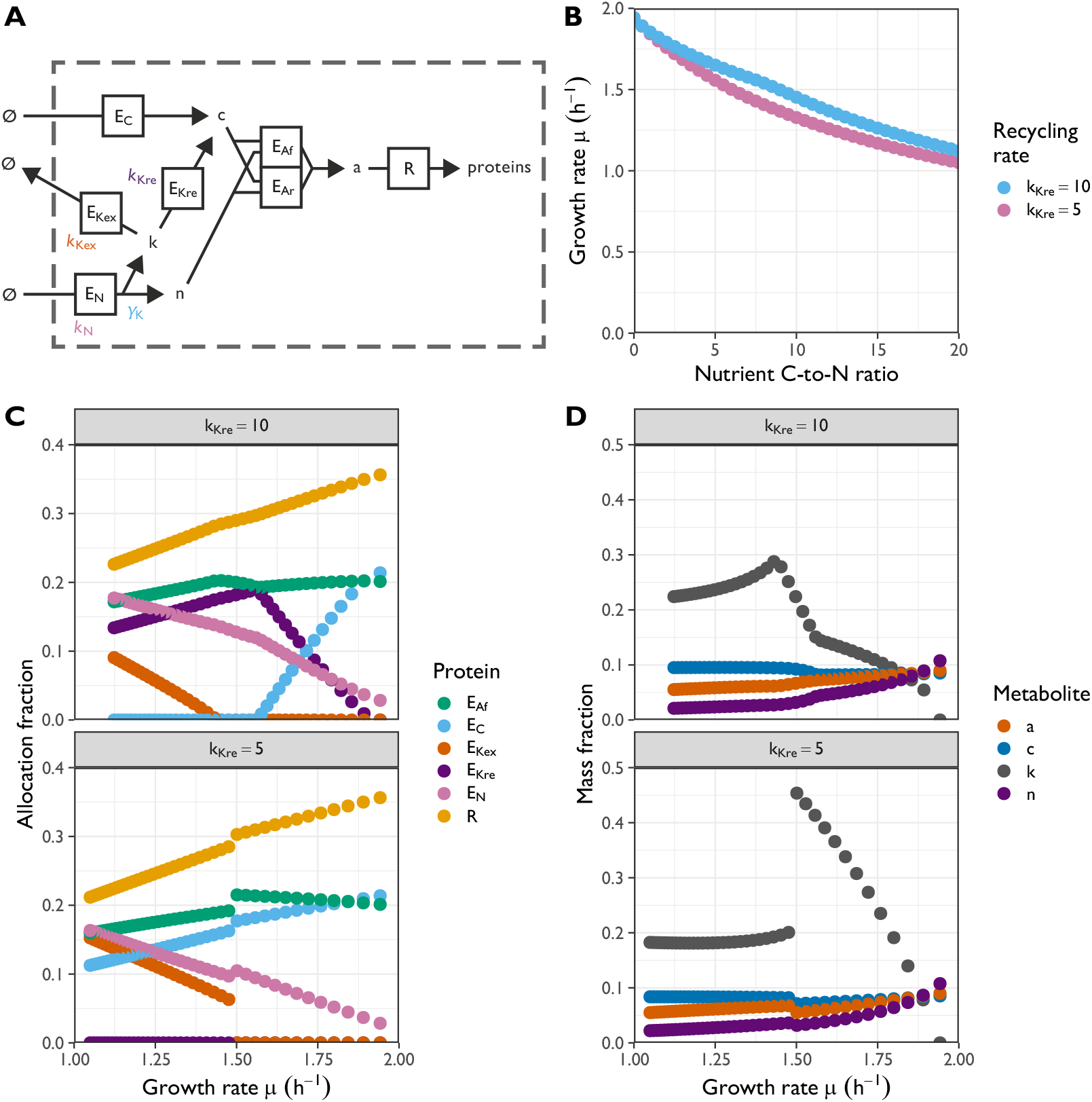
Analysis of the full model, including recycling and excretion of ketoacid under modulation of the carbon-to-nitrogen ratio in the nitrogen source for two choices of the ketoacid recycling rate k_K_re__. The carbon-to-nitrogen ratio was varied between 0 and 20 in steps of 0.5 and the ketoacid stoichiometry γ_K_ was calculated from this as described in the text. Allocation fractions were chosen to maximise the growth rate for each γ_K_ and k_K_re__. **A.** Illustration of the model. **B.** Growth rate μ as a function of the nitrogen source,s carbon-to-nitrogen (C-to-N) ratio for the respective ketoacid recycling rates. **C.** Optimal protein allocation fractions as a function of growth rate μ for the two recycling rates. Purely respiratory enzymes E_Ar_ and housekeeping proteins Z were omitted from the figure to improve clarity; E_Ar_ was not expressed for any of the parametrizations shown. **D.** Steady-state mass fractions of metabolites as a function of growth rate μ.

The behaviour of the model that included ketoacids depended qualitatively and quantitatively on the values of key parameters, particulary on the efficiencies (catalytic rates) of the enzymes involved and on the ketoacid stoichiometry γ_K_. Recycling enzymes were only expressed if their efficiency (catalytic rate) k_K_re__ was large. We first explored the model behaviour for two choices of this parameter, modulating γ_K_ but leaving fixed all other metabolic parameters. The results are shown in Figs 3B-D.

Based on the types of proteins that were expressed, there were three qualitatively different growth regimes for k_K_re__ = 10 and two for k_K_re__ = 5. One important trade-off here is between the cellular cost of expressing ketoacid processing enzymes and the cost of carrying the excess metabolite for cell growth. When recycling enzymes were inefficient and costly (k_K_re__ = 5, bottom panels), but the carbon-to-nitrogen ratio was still below a certain threshold, the ketoacid metabolites built up considerably (up to almost half the total biomass). In this regime, this still lead to faster growth than expressing either recycling or excretion enzymes would have. However, with the amino acid nitrogen source containing relatively more carbon, the growth-optimized cell excreted ketoacids and the internal concentration was approximately stable across the range of growth rates investigated. When recycling enzymes were efficient ((k_K_re__ = 10, top panels), re-cycling replaced the uptake of carbon through the canonical carbon pathway, i.e. no carbon uptake enzyme EC was expressed below a critical growth rate. For both choices of k_K_re__, in the fastest-growing regime, no excretion took place, i.e. the excretion enzyme EKex was not expressed; but in the slowest-growing regime, ketoacid excretion was required to optimize growth. Curiously, we found a recycling-only regime with neither canonical carbon uptake nor ketoacid excretion. In this latter regime, all carbon in the biomass has its origin in the amino acid from the nutrient, and all the carbon from the nutrient made its way into the cell.

We note that across all simulations, no respiratory enzyme EAr was expressed. Furthermore, the optimal allocation to ribosomal proteins deviated discontinuously from the ribosomal growth law for both choices of k_K_re__ when the ketoacid stoichiometry γ_K_ was varied.

### 2.4 Different nutrient environments (parametrisations) induced complex trade-offs between carbon uptake, ketoacid recycling, and excretion

We further explored the five different growth regimes across a more extensive parameter sweep. The ketoacid stoichiometry γ_K_ is related to the ratio of carbon and nitrogen atoms in a nutrient molecule:

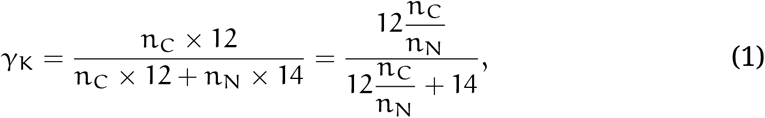

where n_C_ and n_N_ are the numbers of carbon and nitrogen atoms in a nutrient molecule, and the factors 12 and 14 account for the approximate molar masses of these two elements. For example, glycine molecules contain two carbon atoms for each nitrogen atom, which would be represented by 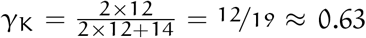, whereas each molecule of isoleucine contains nine carbon atoms, for 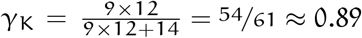.

Alongside the ketoacid stoichiometry γ_K_, we varied the maximal turnover rates (efficiencies) k_N_, k_K_ex__, and k_K_re__ of the enzymes involved in ketoacid metabolism, as plotted in Fig. 4. This showed that the five regimes highlighted in the previous section were universal. The optimally allocated cell expressed only one or two out of the ketoacid recycling, ketoacid excretion, and carbon uptake enzymes depending on the parametrisation (Fig. 3). Neither recycling nor excretion was expressed in nitrogen sources not also containing carbon, and additionally this was optimal even for carbon-containing nitrogen sources when the ketoacid recycling and excretion enzymes were inefficient relative to the nitrogen uptake enzyme (Fig. 4ADE). The value of the ketoacid recycling rate k_K_re__ below which ketoacid recycling was suboptimal depended weakly on the ketoacid stoichiometry (Fig. 4A). When recycling enzymes were expressed, the trade-off between the excretion and ketoacid uptake was heavily influenced by the nitrogen source’s carbon content and all three enzyme rates (Fig. 4ABC). Low-carbon nutrient sources required additional carbon uptake whereas high-carbon nutrient sources generally required ketoacid excretion, with pure recycling being favoured in regimes with intermediate carbon content or inefficient excretion.

**Figure 4:**
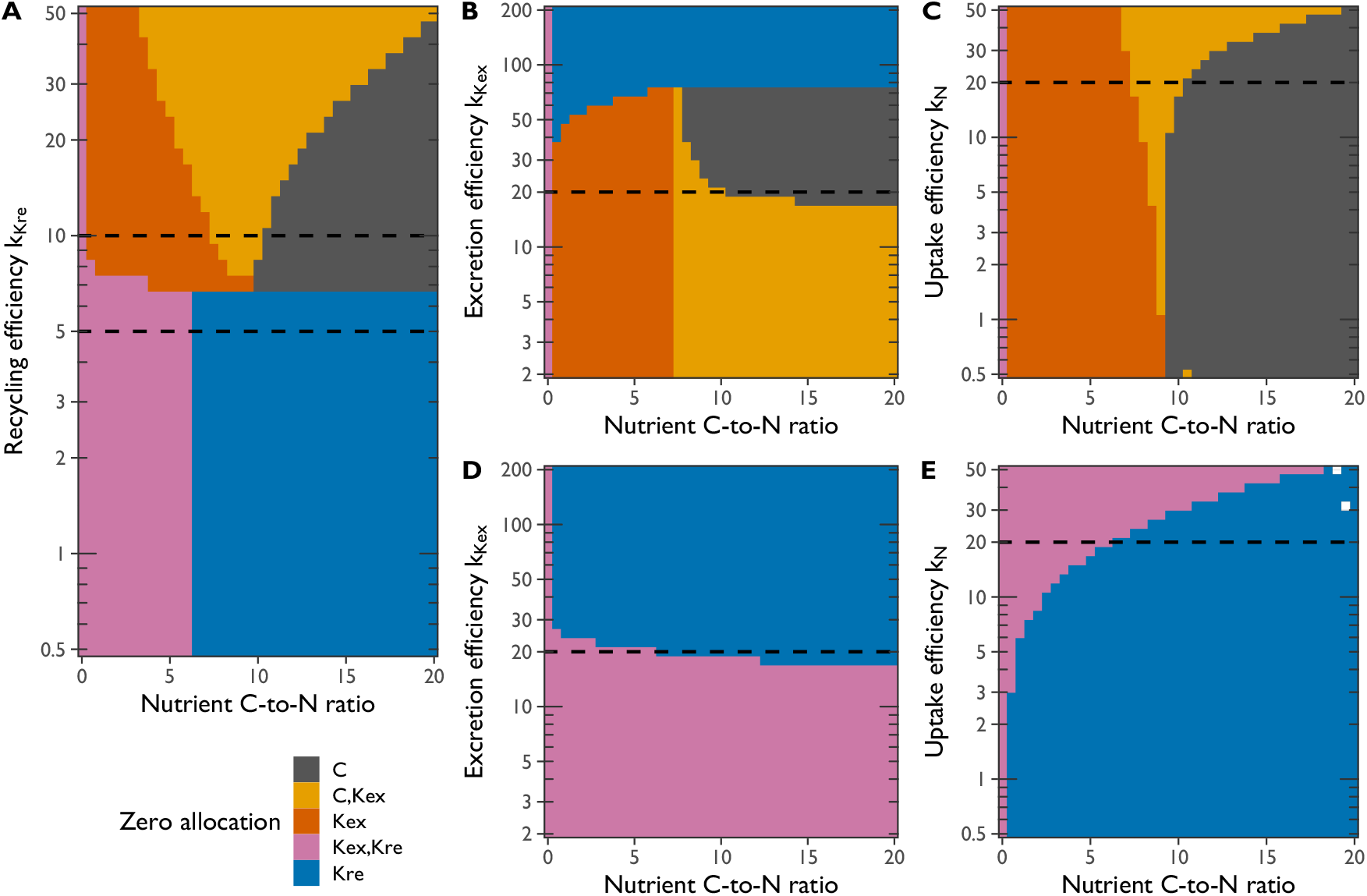
Phase diagrams of ketoacid recycling, excretion, and carbon uptake under perturbations of enzyme efficiencies and nutrient carbon-to-nitrogen ratio. For each figure, the optimal allocation was determined for 41^2^ combinations of the ketoacid stoichiometry γ_K_ and one of the enzyme efficiencies k_K_re__, k_K_ex__, and k_N_. As before, the carbon-to-nitrogen ratio was varied between 0 and 20 in steps of 0.5 and the ketoacid stoichiometry γ_K_ was subsequently calculated. Furthermore, the enzyme efficiencies were chosen to be equidistant after log-transformation. In the figure, colours indicate which of the enzymes were not expressed in the optimal allocation, and dashed lines indicate parameter values that were fixed in the other panels and in 3. **A.** Ketoacid recycling rate k_K_re__ varied, ketoacid excretion rate k_K_ex__ = 20.0 h^-1^ and nutrient uptake rate k_N_ = 20.0 h^-1^ held fixed. **B.** k_K_ex__ varied, k_K_re__ = 10.0 h^-1^ and k_N_ = 20.0 h^-1^ fixed. **C.** k_N_ varied, k_K_re__ = 10.0 h^-1^ and k_K_ex__ = 20.0 h^-1^ fixed. **D.** and **E.** as B. and C., but with k_K_re__ = 5.0 h^-1^.

### 2.5 Approximately optimal allocation towards ribosomes could result from amino-acid regulation

Until now, we have explored the model using defined modulations of one or more rate parameters and the ketoacid stoichiometry. Across all of these, we found that the ribosomal growth law (ribosomal proteome allocation fraction f_R_ ∝ μ) was robustly satisfied. Moreover, the amino acid abundance a appeared to be similarly correlated with the growth rate. We wondered if this was an artefact of our parametrization approach or a deeper property of the model and used a random sampling parametrization strategy to further study this behaviour. We sampled 100 triplets of the rate parameters (k_C_, k_N_, k_A_f__) from independent uniform distributions with support [0, 20], set k_A_r__ = 0.5k_A_f__, and matched these samples to four choices of the ketoacid stoichiometry γ_K_, representing carbon-to-nitrogen ratios of 0, 3, 6, and 12, and two choices of the ketoacid recycling efficiency k_K_re__.

In the ketoacid-free model (γ_K_ = 0) with random rate parameters, the ribosomal growth law was nearly exactly satisfied (Fig. 5A), even though expression of the two transporters and the metabolic enzyme was highly variable between parameter choices. Furthermore, the concentration of the amino acid a was closely related to growth rate as well (Fig. 5B). It follows from the growth rate correlations for both f_R_ and a that the two were correlated themselves as well (Fig. 5C). A linear fit f_R_ ∝ (a – a_0_) closely approximated the observed relation.

**Figure 5:**
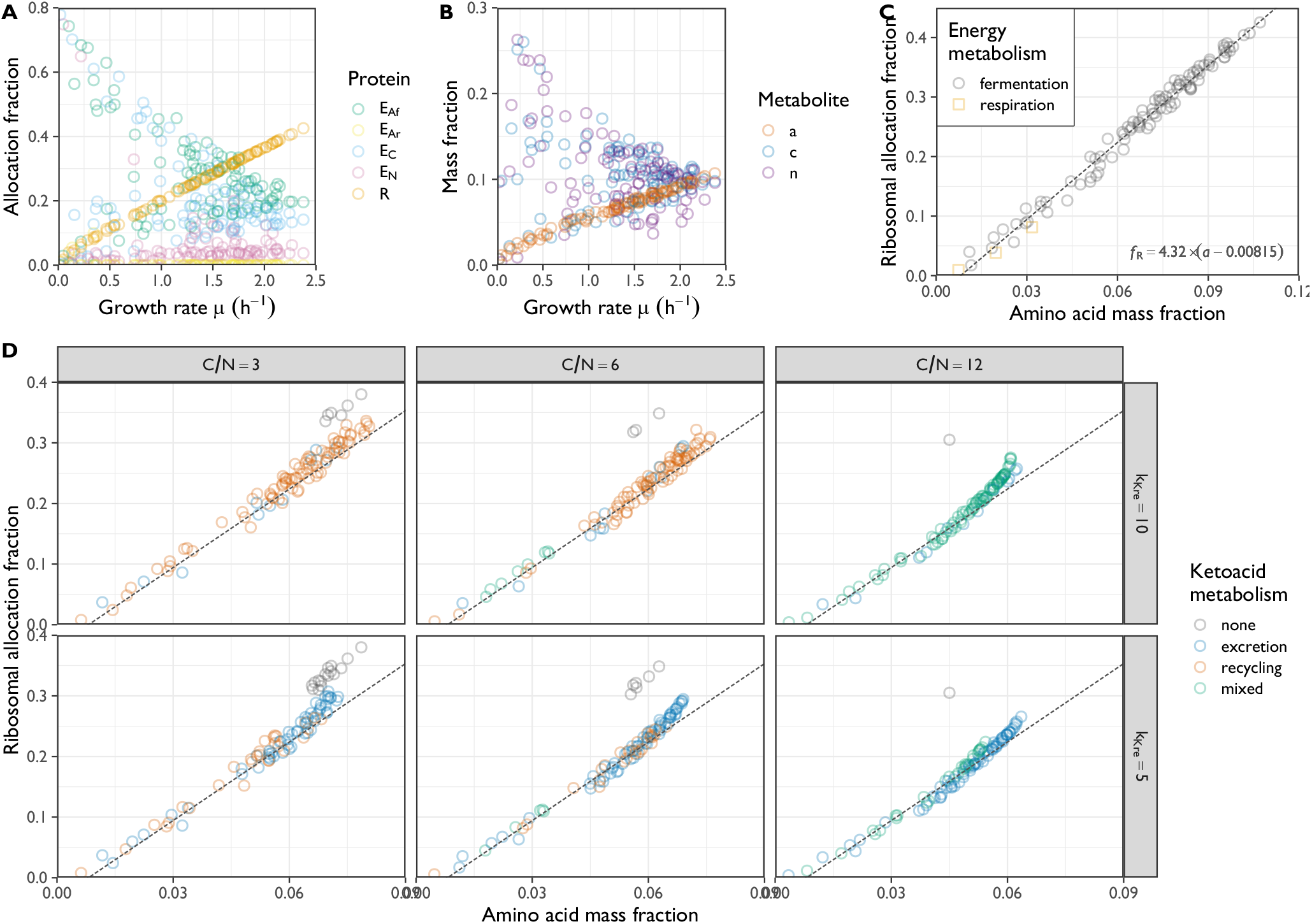
Analysis of the full model with randomly chosen rate parameters, exploring the relation between amino acid precursor abundance and ribosomal allocation. The enzyme efficiencies k_A_f__, k_C_, and k_N_ were randomly drawn from separate uniform distributions ranging between 0 and 20.0 h^-1^, and the respiratory enzyme efficiency k_A_r__ was set to 0.5k_A_f__. The ketoacid recycling efficiency k_K_re__ and carbon-to-nitrogen ratio were initially fixed to 10.0 h^-1^ and 0 respectively (for A.-C.), and then varied jointly (for D.). The allocation fractions were chosen to maximise the growth rate for each set of parameters. **A.** Optimal protein allocation fractions as a function of the growth rate μ. Housekeeping proteins and ketoacid-processing enzymes were omitted to improve clarity. **B.** Steady-state biomass fractions of amino acid a, carbon a, and free nitrogen n plotted against the growth rate μ. **C.** Scatter plot of the amino acid concentrations a and ribosomal allocation fractions f_R_, indicating the type of energy metabolism that optimised growth (fermentation: only E_Af_ expressed; respiration: only E_Ar_ expressed). The dashed line represents an ordinary least squares linear model fit to the data. **D.** As C. for three nonzero choices of the carbon-to-nitrogen ratio (C/N) and two choices of the ketoacid recycling efficiency k_K_re__. Points are coloured according to whether none, one, or both of the ketoacid excretion enzyme E_Kex_ and ketoacid recycling enzyme E_Kre_ was expressed. The dashed line indicates the fit from panel C. (no further fitting was performed here).

We further explored the relation between f_R_ and a in the presence of complex nitrogen sources such as amino acids, when additionally the recycling and/or excretion of ketoacid is accounted for (Fig. 5D). For large carbon-to-nitrogen ratios, the relation between f_R_ and a may be better described by a nonlinear relation, al-though the fit for f_R_ ∝ (a – a_0_) obtained for nitrogen sources without carbon was still a close approximation. Only when the optimal solution involved ketoacids building up in the cell without recycling or excretion did the linear fit break down entirely. Since our model does not account for the toxicity of the intermediate ketoacids beyond their passive drag on growth, this situation is not likely to occur in real cells.

All together with the previous sections, these results suggest that a single linear regulation of ribosomes by amino acids can achieve near optimal allocation of ribosomes. For the regimes explored here, a simple proportionality f_R_ = δa such as proposed by Bertaux et al. (2020) would work reasonably well, though our introduction of an offset a_0_ (i.e. f_R_ = δ(a – a_0_)) improved the fit. Interestingly, if f_R_ ∝ a is chosen and the ribosomes are implemented with Michaelis–Menten kinetics the ribosomal growth law follows analytically. A derivation of this statement is presented in Supp. Text 5.4; it holds whether or not there is an offset a_0_ in the f_R_–a relationship. We note that we did not observe a simple relationship between the allocation of other non-ribosomal proteins and the metabolite concentrations.

## 3 Discussion

A holistic understanding of growth, gene expression, and resource allocation can be codified in and achieved by coarse-grained resource allocation models (C-GRAMs). Here, we constructed a minimal C-GRAM of microbial metabolism that accounted for (i) the metabolism of both carbon and nitrogen, (ii) the different proteomic and kinetic efficiencies of respirofermentative and respiratory energy metabolism, and (iii) the stoichiometry of complex nitrogen sources that contain a carbon backbone, such as amino acids. We optimized the allocation towards different protein classes so as to maximise the growth rate.

Different parameter sweeps of the model allowed us to explore different nutrient environments. The optimal allocation and resulting growth rate co-varied according to strikingly regular behaviour. In particular, the Monod law and ribosomal growth law were very robust. Furthermore, we described growth on carbon-containing nitrogen sources by introducing the possibility of recycling and/or excreting residual carbon, which induced complex trade-offs in metabolism. Notably, both the optimal allocation towards ribosomes and their substrate (internal amino acids) varied approximately linearly with the growth rate, leading to an also approximately linear relationship between the two.

### 3.1 Protein reserves and simple feedback of free amino acids setting ribosome allocation

Our assumption that expression of all proteins was optimised for growth in any given condition considerably simplified the model parameterization. Precise optimality conditions have been formulated for the *E. coli* carbon uptake system and gene expression indeed maximised the growth rate for several carbon sources (Towbin et al. 2017). Furthermore, a recent theoretical advance pointed out a general method of adapting gene expression control towards the optimum (Planqué et al. 2018), However, recent evidence has challenged the view that all allocation is growth-optimal. It is thought that significant fractions of the *E. coli* (Valgepea et al. 2013; Peebo et al. 2015; Mori et al. 2017) and budding yeast (Metzl-Raz et al. 2017; Yu et al. 2020) proteome are not necessary for sustaining the growth rate. In particular, central carbon metabolism has been suggested to have a large reserve capacity, suggesting that many enzymes are not utilised solely to maximise metabolic fluxes (O’Brien et al. 2016; Christodoulou et al. 2018; Yu et al. 2021). Pools of proteins held in reserve may be beneficial instead by their ability to support adaptation to environmental changes.

An intermediate step between a fully growth-optimised and a dynamically regulated allocation model may be the proportional regulation fR ∝ a. We showed this to be a good approximation to the growth-rate-maximising allocation; additionally, it was robust to many variations of the parameters describing the nutrient quality. The simplicity of this relation is remarkable: in principle, optimal allocation could depend on all internal metabolite concentrations and be highly nonlinear, instead of this linear dependence on only a single metabolite. Explicit regulation of bacterial ribosome synthesis mediated by a single metabolite, guanosine tetraphosphate (ppGpp), was thoroughly explored in a coarse-grained model by Bosdriesz et al. (2015). Recently, ppGpp was shown to regulate the growth rate and ribosome content in *E. coli* by sensing the instantaneous translation elongation rate (Wu et al. 2022).

Notably, neither our growth-optimised model nor one implementing ribosomal allocation proportional to internal amino acids explicitly account for a reserve pool of ribosomes not actively involved in translation. Such a pool has been held responsible for the observed offset φ_R0_ in the ribosomal growth law φ_R_ = φ_R0_ + σ^-1^ μ (Dai et al. 2016; Metzl-Raz et al. 2017; Mori et al. 2017). Although our model did not implement any inactive ribosomes, we still observed an offset greater than zero. We explain this effect by the fact that the ribosomes in our model are not fully saturated with substrate, even when growth is optimised. Specifically, linearly correlated ribosome allocation and amino-acid abundance (either through optimisation or explicit regulation), combined with nonlinear Michaelis–Menten ribosome kinetics, resulted in a offset

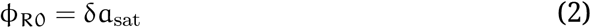

when f_R_ = δ(a – a_0_) and 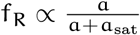. Naturally, ribosome saturation and inactivation are not mutually exclusive, and a quantitative understanding of their relative importance will have profound implications on our understanding of the interplay between ribosome synthesis, excess translational capacity, and cell growth (Dai and Zhu 2020).

### 3.2 The fate of the carbon backbone

We next discuss the fate of the carbon backbone from complex nitrogen sources, as modelled first in the model presented here. In our model, modulations of the ketoacid stoichiometry γ_K_ gave rise to a wide range of growth rates in a monotonically decreasing manner. However, a nitrogen source,s quality is not solely determined by its carbon-to-nitrogen ratio. For example, glycine and tryptophan media gave rise to very similar growth rates in *S. pombe* (Kleijn et al. (2022), but carbon is present in a 2:1 ratio in glycine and in a 5.5:1 ratio in tryptophan (see Supp Text, Table 4). This strongly suggests that each growth medium is not only associated with γ_K_, but that at least one out of the enzymatic rates k_N_, k_K_ex__, and k_K_re__ must also be modulated by the choice of nitrogen source. While our model provides a framework for understanding the optimality of recycling and excretion of carbon, we refrained from a full parameterization of growth on specific nitrogen sources. Unlike translation and central carbon metabolism, the topology of amino acid uptake pathways is rather poorly conserved between organisms. In practice, then, whether recycling or excretion is preferred for a given amino acid nitrogen source depends on which reactions are available to the organism, and how efficient they are. If ketoacid recycling effectively feeds into other synthesis pathways, this would correspond to a large k_K_re__ in the model; on the other hand, the effectiveness (or absence) of suitable excretion pathways would influence the value of k_K_ex__.

**Table 1:** Variables (metabolites and proteins) implemented in the full model.

| Type | Description | Concentration | Allocation fraction |
| --- | --- | --- | --- |
| Metabolite | Ketoacid | $k$ | N/A |
| | Carbon | $c$ | |
| | Amino acid | $a$ | |
| | Free nitrogen | $n$ | |
| Protein | Ketoacid recycler | $e_{K_{re}}$ | $f_{K_{re}}$ |
| | Ketoacid excreter | $e_{K_{ex}}$ | $f_{K_{ex}}$ |
| | Carbon uptake | $e_C$ | $f_C$ |
| | Respirofermentative enzyme | $e_{A_f}$ | $f_{A_f}$ |
| | Purely respiratory enzyme | $e_{A_r}$ | $f_{A_r}$ |
| | Nitrogen-compound uptake | $e_N$ | $f_N$ |
| | Ribosome | $r$ | $f_R$ |
| | Housekeeping (non-metabolic) | $z$ | $f_Z$ |

**Table 2:** Parameters (Michaelis constants, enzyme efficiencies, and stoichiometries) implemented in the full model.

| Type | Description | Symbol | Default value |
| --- | --- | --- | --- |
| Michaelis constant | Ketoacid | $k_{sat}$ | 0.0167 |
| | Carbon | $c_{sat}$ | 0.0167 |
| | Amino acid | $a_{sat}$ | 0.0167 |
| | Free nitrogen | $n_{sat}$ | 0.0167 |
| Enzyme efficiency | Ketoacid recycler | $k_{K_{re}}$ | $10.0 \text{ h}^{-1}$ |
| | Ketoacid excreter | $k_{K_{ex}}$ | $20.0 \text{ h}^{-1}$ |
| | Carbon uptake | $k_C$ | $10.0 \text{ h}^{-1}$ |
| | Respirofermentative enzyme | $k_{A_f}$ | $15.0 \text{ h}^{-1}$ |
| | Purely respiratory enzyme | $k_{A_r}$ | $7.5 \text{ h}^{-1}$ |
| | Nitrogen-compound uptake | $k_N$ | $20.0 \text{ h}^{-1}$ |
| | Ribosome | $k_R$ | $6.46 \text{ h}^{-1}$ |
| Mass stoichiometry | Carbon consumed per amino acid produced in respirofermentative metabolism | $\alpha_{C_f}$ | 48/55 |
| | Carbon consumed per amino acid produced in purely respiratory metabolism | $\alpha_{C_r}$ | 24/31 |
| | Ketoacid produced per nitrogen compound consumed | $\gamma_K$ | 0 |

**Table 3:** Common choices for the abundance measure in whole-cell models. Note that molar concentrations 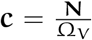 and mass concentrations 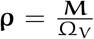 are composite measures.

| absolute abundance $X$ | system size $\Omega$ | relative abundance $x$ |
| --- | --- | --- |
| molecule number $N$ | total number of molecules $\Omega_N$ | molar fraction |
| mass $M$ | cell mass $\Omega_M$ | mass fraction $\phi$ |
| occupied volume | cell volume $\Omega_V$ | volume fraction |

**Table 4:** Carbon-to-nitrogen ratios of selected amino acids. The amino acids are ordered according to the growth rate that they support in *S. pombe* minimal media (Kleijn et al. 2022). Note that tryptophan molecule contains two nitrogen atoms; the other selected amino acid molecules contain no nitrogen apart from the backbone amino group. A natural choice for the ketoacid stoichiometry parameter γ_K_ from the model is given by 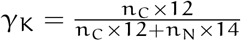.

| Amino acid | C:N ratio | $n_C$ | $n_N$ | $\gamma_K$ |
| --- | --- | --- | --- | --- |
| Tryptophan | 5.5 | 11 | 2 | $61/80 \approx 0.83$ |
| Glycine | 2 | 2 | 1 | $12/19 \approx 0.63$ |
| Phenylalanine | 9 | 9 | 1 | $54/61 \approx 0.89$ |
| Serine | 3 | 3 | 1 | $36/50 = 0.72$ |
| Isoleucine | 6 | 6 | 1 | $36/43 \approx 0.84$ |
| Proline | 5 | 5 | 1 | $30/37 \approx 0.81$ |
| Glutamate | 5 | 5 | 1 | $30/37 \approx 0.81$ |

It is challenging to condense whole-genome metabolic models into C-GRAMs due to the complexity of metabolism; the presence of many parallel pathways, moonlighting enzymes, and metabolic cycles are just three examples. However, some progress may be made in adapting our generic C-GRAM to specific organisms. The metabolism of indigestible ketoacids has been well studied in *S. cerevisiae*, whose excretions of such “fusel oils” can spoil industrial applications (Godard et al. 2007). Recently, high-quality metabolic maps that are aware of limited resource allocation in multiple cellular compartments have been developed for *S. cerevisiae* and *S. pombe* (Elsemman et al. 2022; Grigaitis et al. 2022). Such models may enable quantitative mappings between the coarse-grained proteome sectors and proteome data in the future.

### 3.3 Explicit overflow metabolism and energy generation

We accounted for the distinction between purely respiratory and respirofermentative growth by adjusting parameters of the enzyme representing the pathway, similar to one approach by Molenaar et al. (2009). Specifically, we adjusted α_C_, which represents the stoichiometry of carbon required for biomass production relative to nitrogen, and k_E_, the efficiency (maximal specific flux) of the pathway. This induced differing behaviours under nitrogen and carbon limitation: carbon limitation induced a switch to fully respiratory growth upon decreasing growth rates but nitrogen limitation did not. However, an important features of fermentation, namely the excretion of overflow metabolites, was not explicitly modelled. Furthermore, we did not include the generation of cellular energy in the form of ATP. These two omissions are related, as ATP is generated in different amounts by fermentative and respiratory pathways. A natural extension of the model would therefore be the addition of an explicit fermentative pathway. Molenaar et al. (2009) showed that this approach may induce mixed expression of the two pathways in a C-GRAM with linear metabolism lacking nitrogen uptake. It is important to consider that in our model, the pathway represented by E_Af_ includes respiration as well as fermentation, and respiratory enzymes and the citric acid cycle are required for amino acid biosynthesis (Malecki et al. 2020).

### 3.4 Non-protein biomass

A second caveat to our choice modelling biosynthesis in one simple pathway is the following. While the parameter α_C_ represented the biomass carbon-to-nitrogen ratio in the model, it cannot be directly equated to observed dry mass compositions in real cells (some examples are collated in Supp Text, Table 5). This is partially due to the inclusion of extra carbon that real cells excrete during fermentation (see previous paragraph and Supp Text 4.3), but also because the biomass of real cells consists of additional macromolecules besides proteins. For example, nucleotides are important cellular constituents with a higher nitrogen content than proteins, and an efficient cell must synthesise them in proportion to ribosomal proteins, as most RNA is ribosomal. Upstream, nucleotide synthesis depends on the pentose phosphate pathway (PPP), which shares multiple intermediate metabolites with glycolysis and amino acid metabolism. The trade-off between glycolytic and PPP flux has been implied to influence the relationships between growth rate, yield, and oxidative stress (Christodoulou et al. 2018; Cheng et al. 2019; Hurbain et al. 2022). A coarse-grained model that includes both protein and nucleotide synthesis must account for the coordination between the two carbon metabolic pathways (Chubukov et al. 2014), considerably increasing its complexity versus the model presented in this study.

**Table 5:** Approximate relative dry mass composition of selected organisms. Most of these numbers were obtained via the Bionumbers database and corresponding identifiers (BNIDs) are indicated (Milo et al. 2010). The protein content of *S. pombe* was approximated from the following observations: (i) the cell measures approximately 14 by 3.5 microns (Fantes and Nurse 1977), resulting in a volume of 0.123 pl if its shape is assumed to be a spherocylinder; (ii) the average dry mass density is 282 pg/pl (Odermatt et al. 2021), such that the dry mass per cell is approximately 35 pg; (iii) the protein content (of haploid wild type cells in glucose minimal medium) is 15 pg/cell (Fantes and Nurse 1977).

| Species | Chemical formula | Protein content (g/g) |
| --- | --- | --- |
| *E. coli* | $CH_{1.77}O_{0.49}N_{0.24}$ (BNID 101800; von Stockar and Liu 1999) | 0.55 (BNID 101436; Neidhardt et al. 1990) |
| *S. cerevisiae* | $CH_{1.613}O_{0.557}N_{0.158}$ (BNID 101801; von Stockar and Liu 1999) | 0.53 (BNID 102328; Ertugay and Hamamci 1997) |
| *S. pombe* | $CH_{1.9}O_{0.61}N_{0.155}$ (de Queiroz et al. 1993) | $\sim 0.43$ |

Another large contribution to the biomass of microbes comes from their cell surface, which mostly consists of carbon. The interplay between cell surface biosynthesis, size homeostasis, and growth has been explored in coarse-grained models of bacteria (Harris and Theriot 2016; Ojkic et al. 2019; Oldewurtel et al. 2019). The relative importance of the cell surface changes with the size and shape of the cell, both of which depend on growth conditions and fluctuate during the cell cycle. Therefore, the dry mass density varies as cell mass and cell volume evolve differently. While balanced growth may be defined in terms of repeated cell cycles and the dynamic equations may be studied outside of steady state, the fluctuating dry mass density invalidates a focus on relative abundances. Accounting for the contribution of the cell surface to biomass in our model would therefore require reformulating the model into absolute abundances and including the effect of the cell cycle, which was outside the scope of this study. However, we expect that the improved understanding of concepts such as the optimisation of resource allocation, the kinetic modelling of coarse-grained pathways, and the parameterization of stoichiometries, as presented in this work, should be useful in the development of surface-aware C-GRAMs.

### 3.5 Conclusion

In summary, we have presented a modelling framework that describes uptake and metabolism of carbon and nitrogen in unicellular microbes in a coarse-grained manner. While the parameterization of the model presented here was chosen to facilitate comparisons with earlier models of *E. coli*, the structure of the model was deliberately kept general. The framework may therefore be applied to study the optimality of gene expression and growth across the tree of life. We hope that extensions of our model will be constructed to describe overflow metabolism, ATP, nucleotide metabolism, a cell surface, and/or cell density fluctuations.

## Acknowledgements

We would like to thank Philipp Thomas, Nick Jones, and Peter Swain for discussions related to some of the material presented here.

## 4 Methods

### 4.1 Dynamic equations describing the metabolic model

The model from Bertaux et al. (2020) served as our starting point. We disregarded non-metabolic proteins and the inhibition of ribosomes, which were both present in the original model, because here we aimed to describe the steady-state behaviour of unperturbed wild-type cultures. We then introduced additional metabolic pathways representing (i) the uptake of carbon and nitrogen, (ii) fermentative and respiratory energy generation, represented by different parametrisations of a similar enzyme, and (iii) the recycling and excretion of carbon from complex nitrogen sources containing both nitrogen and carbon. Simplifications of the model were obtained by setting some model parameters to zero.

The reactions included in the full model are pictured in 3A; the interpretations of the variables and parameters in the model are given in 1 and 2. These reactions were modelled by the following formalism of ordinary differential equations (ODEs), which is explored in more detail in 5.2. The time evolution of the concentration vector *x* was decomposed and given by:

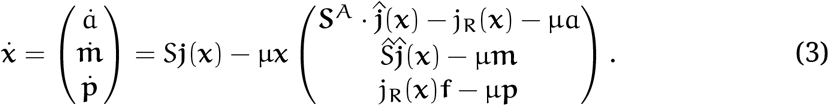

The concentrations described by this equation are those representing the amino acids a, the other metabolites **m** = (k, c,n), and the proteins **p** = (e_K_re__, e_K_ex__, e_C_, e_A_f__, e_A_r__, e_N_, r, z). The ribosomal allocation is given by the vector **f** = (f_K_re__, f_K_ex__, f_C_, f_A_f__, f_A_r__, f_N_, f_R_, f_Z_). Fluxes catalysed by the proteins are given by the vector

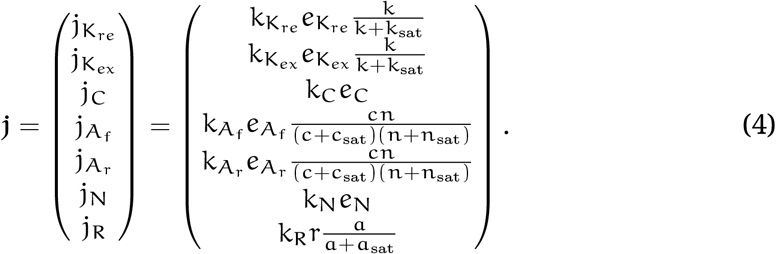

The enzymes were chosen to follow Michaelis–Menten kinetics (for single-substrate enzymes) and products thereof, such that the rate laws were linear in the enzyme concentration, linear in the substrate concentrations at low concentrations, and saturated at high substrate concentrations. Details on the enzyme kinetics are provided in Supp. Text 5.3. Note that the housekeeping proteins (with concentration z) do not catalyse any enzymatic reaction and therefore did not feature in the fluxes. Finally, the stoichiometry matrix was given by

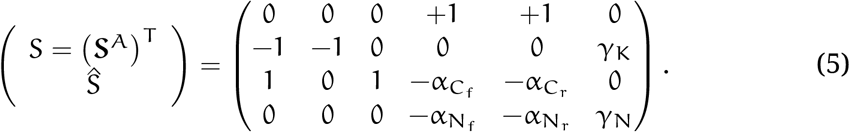

Mass balance was maintained in all internal reactions, meaning columns in the stoichiometry matrix representing these totaled zero. However, the carbon and nitrogen transporters imported nutrients from the environment (so columns totaled +1), and the ketoacids could be excreted to the environment by the respective enzyme (so this column totaled −1). These considerations imposed the following constraints:

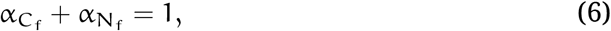

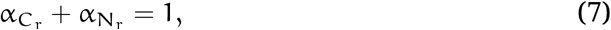

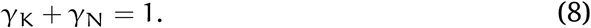

Using the above definitions, the ODEs describing the system can be written as:

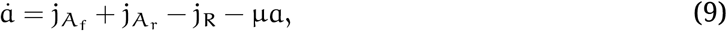

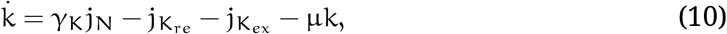

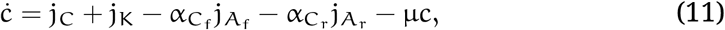

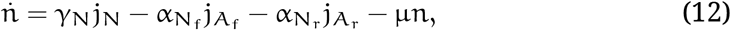

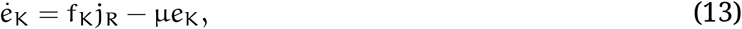

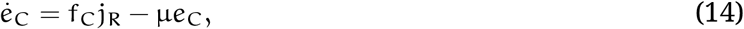

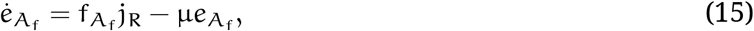

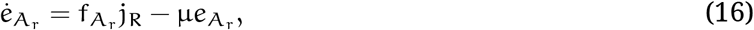

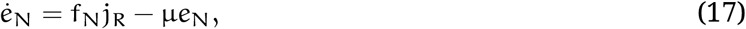

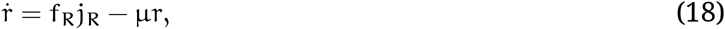

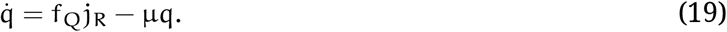

The growth rate μ is found by taking the sum of these equations. Furthermore, using the allocation constraint ∑_i_ f_i_ = 1 and the concentration constraint ∑_i_ *x*_i_ = 1, such that ∑_i_ *ẋ*_i_ = 0, gives

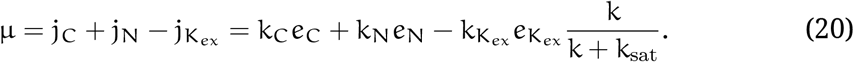

This is equal to the net import of nutrient from the environment (uptake minus excretion).

### 4.2 Balanced growth

As mentioned in the main text, we aimed to describe gene expression allocation under balanced growth in defined environments. The state of balanced growth was represented in the model by the steady-state of equation (3). To compute the steady-state numerically, we evolved these equations until a steady-state was reached.

Given a model parameterization (see 4.3), an initial condition was partially guessed from the allocation fractions. This used that the protein content in steady-state is proportional to the proteome allocation. However, the initial metabolite fraction was established manually as 0.05 for the relative concentration of each metabolite. In other words, the initial concentration vector *x_0_* was set to

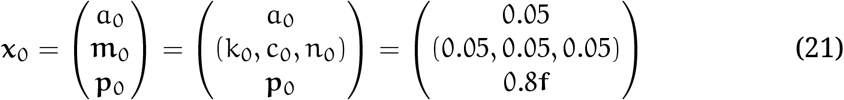

with f the given allocation vector.

The simulations were implemented in the Julia programming language, using the DifferentialEquations.jl ecosystem (Rackauckas and Nie 2017). For solving the concentration ODEs towards steady-state, we used the Rodas5 solver.

### 4.3 Parameterization

Unicellular organisms grow in one of two principal states: respirofermentative growth, and entirely respiratory growth. From the point of view of coarse-grained modelling, there are two primary differences between the two. On the one hand, the fermentation pathway consists of few different enzymes. Although the individual enzymes are highly abundant, they are highly efficient. The aggregate effect on the total expression burden is that the fermentation enzymes comprise a markedly smaller proteome fraction than the respiratory pathway would when providing equal biomass and energy production. On the other hand, fermentation requires more nutrients from the environment: carbon is consumed rapidly and converted into ethanol or acetate. We encoded the efficiency in the rate parameter k_*A*_, and the carbon usage in the parameter α_*C*_. Specifically, with respirofermentative growth represented by the subscript f and purely respiratory growth by the subscript r the above considerations require k_A_f__ > k_A_r__ and α_C_f__ > α_C_r__.

To obtain a rough estimate of the stoichiometry parameter in respiratory growth α_C_r__, we used that proteinogenic amino acids contain approximately 4 carbon atoms per nitrogen atom. Using molar masses of 12 g/mol and 14 g/mol for carbon and nitrogen, respectively, this ratio translated to (mass-action)

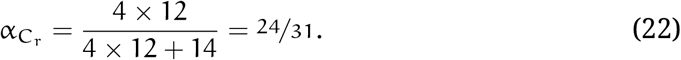

A further rough estimate, namely that the excess carbon excreted in fermentation is approximately equal in amount to the carbon converted to biomass, gave the stoichiometry in respirofermentative growth as

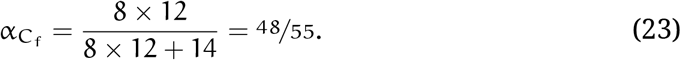

The full metabolic model, described in 4.1, was explored in Figures 3, 4, and 5. The simplified models described earlier were obtained by setting key parameters to zero. For the model with parallel carbon and nitrogen assimilation pathways, the lack of ketoacid recycling or excretion (Figure 2) was represented by the ketoacid stoichiometry parameter γ_K_ = 0. Additionally, the initial ketoacid fraction k_0_ was set to zero, and no allocation was made to the ketoacid enzymes, i.e. f_K_ex__ = f_K_re__ = 0. For the initial core model, which did not distinguish respirofermentative and respiratory growth, f_*A*_r__ = 0 was additionally forced.

### 4.4 Optimising ribosomal allocation to maximise steady-state growth rate

We explored the behaviour of the model under parameter modulations that represented different growth environments, while assuming that growth was optimized to suit this environment. The allocation vector **f** from equation (3) was therefore chosen not as a free parameter, but rather as the result of an optimisation routine that maximised the growth rate μ. The fraction of housekeeping proteins f_Z_ was excluded from the optimisation. The optimisation problem can be defined as finding the allocation fraction **f** that maximises the growth rate μ, while satisfying the allocation constraint

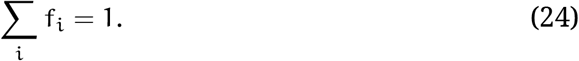

The computation of the optimal allocation fraction is complicated, because the function μ(**f**) is only defined implicitly: a steady-state must be found for each choice of **f** before the steady-state growth rate can be extracted. The nonlinearity of the Michaelis–Menten kinetics prohibits an explicit solution for μ in terms of **f**, which would be required by gradient-based based optimisation routines. We were therefore restricted to using a gradient-free optimisation routine, for which we chose the Nelder–Mead algorithm as defined in the Optim.jl package (Nelder and Mead 1965; Gao and Han 2012; Mogensen and Riseth 2018). We used tolerances of 1.0 × 10^-10^ and set the initial simplex to `Optim.AffineSimplexer(a = 0.0, b = −0.1)`. The optimization objective was set to minimize the doubling time 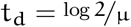.

The allocation constraint (24) further complicated the optimisation procedure. We eliminated one element of the allocation vector by imposing the allocation constraint. However, the Nelder-Mead sampling strategy still allowed for situations where this constrained element became negative. As a practical solution for such situations, and for allocation fractions whose sum exceeded the constraint, we evaluated the doubling time as the inverse of the machine precision, which served as a predefined very large value.

The Nelder–Mead algorithm further proved to be sensitive to numerical inaccuracies when the optimal allocation fraction had zero elements, i.e. the optimal cell entirely lacked expression of some proteins. Therefore, we ran the optimisation procedure several times, each with different allocation fractions set to zero and excluded from the optimisation. Each iteration resulted in one optimal allocation vector; the one with the largest growth rate (smallest doubling time) was selected as the global optimum.

## 5 Supplementary Text

### 5.1 Earlier coarse-grained modelling of unicellular organisms

#### 5.1.1 Summaries of five coarse-grained resource allocation models

As explained in the main text, we built on and extended earlier coarse-grained resource allocation models (C-GRAMs). To highlight the connections between and similarities of the existing approaches, and to facilitate a comparison with the model presented here, we summarise the approach taken by five earlier models.

First, Maitra and Dill (2015) formulated a minimal model of cells with a single variable representing global energy levels, and dividing biomass between metabolic enzymes and ribosomal proteins. The same authors later extended this to include the effect of ribosomal inhibitors (Maitra and Dill 2016). They considered the energy efficiency (growth rate divided by energy production flux) in steady-state growth, which they showed to depend on the allocation of ribosomes; the optimal efficiencies were close to known experimental observations. It was assumed that ribosomal proteins were stable, but non-ribosomal proteins were degraded; with this assumption the nonzero intercept of the ribosomal growth law was proportional to the degradation rate. Under slow growth, the cell spent many resources replenishing proteins that were being degraded, but the efficiency in-creased with the growth rate as increasing levels of external sugar lead to a larger allocation towards ribosomal proteins.

Next, Weiße et al. (2015) constructed a model incorporating three important trade-offs in cellular growth, coupling gene expression to growth. The trade-offs considered were incorporated by means of finite pools of (i) intracellular energy, utilised in all biochemical processes contained in the model; (ii) ribosomes, which produced all proteins; and (iii) cell mass, meaning that production of one type of protein came with an opportunity cost. This model included both transcription and translation, and the availability of intracellular energy levels was assumed to determine transcription and translation rates. Most parameters were taken from the *E. coli* literature, except for a handful of gene expression parameters. These came from a Bayesian (Markov-chain Monte Carlo) fit to the ribosomal growth law data by Scott et al. (2010). The model provided a good balance of interpretability and detail, allowing the authors to explain gene dosage compensation, the interplay between a synthetic circuit and its host cell dynamics, and the evolutionary stability of metabolic gene expression regulation. One notable inter-pretation of this model was that of the ATP-equivalent intracellular energy variable. This was consumed by translation and was therefore alike to amino acids. Furthermore, a good fit to the ribosomal growth law required ribosomal mRNA transcription to be induced at energy levels two orders of magnitude less than non-ribosomal mRNA.

A detailed but still coarse-grained model of resource allocation, growth, and gene expression in bacteria was developed by Liao et al. (2017). They described the dynamics of three protein species, including ribosomal proteins, with their corresponding transcripts, as well as rRNA and tRNA, ATP and free amino acid levels, RNA polymerase levels, and the abundance of the master regulator ppGpp. The model was used to investigate the burden of synthetic constructs, provided a single-cell description of a noncooperative positive feedback loop, and the dynamics of a toggle switch at single-cell, population, and ecological scales. The authors used a combination of widely available data, parameters inferred in earlier studies, and additional Bayesian parameter inference to infer all of 97 parameters.

As seen by this example, complex models accounting for many biological processes also require many parameters, whose inference is nontrivial. This is especially important for research into organisms other than *E. coli*, because fewer experimental results are available. In contrast, a more minimal approach was pioneered by Molenaar et al. (2009), who limited themselves to describing protein abundances and considered the proteome allocation that maximised the growth rate instead of implementing explicit regulatory processes. They described a two-step carbon catabolism with two parameterizations representing metabolically and catabolically efficient metabolism, nitrogen uptake, and ribosomal consumption of both to supply protein production. Molenaar et al. (2009) showed that both growth-rate-correlated gene expression and discontinuous switches between metabolic strategies at specific growth rates could result from simple optimisation of ribosomal allocation for maximal growth rate.

Such a minimal, protein-only, approach was also taken in our group by Bertaux et al. (2020), and the C-GRAM described in this paper is a further development of that model. It introduced explicit cell cycle proteins to describe the dependence of cell size as a function of growth rate under three types of modulation: nutrient limitation, expression of useless proteins, and translational inhibition. One innovation was the inclusion of metabolites in the system size. Furthermore, unlike the growth-rate maximization employed by Molenaar et al. (2009), Bertaux et al. (2020) considered a simple regulation of ribosomal allocation as proportional to their substrate and showed that this strategy obtained near-optimal growth rates.

#### 5.1.2 Consensus model behaviours and important differences

A consensus result of all these models is a hyperbolic dependence of the growth rate on the concentration of external nutrients. This is called the Monod law and it is one of the first global experimental observations of bacterial cultures (Monod 1949). The models also all reproduce the ribosomal growth law, and in fact both the Monod law and the ribosomal growth law have been used to quantitatively fit the model parameters. As mentioned above, the allocation to ribosomes contains a sizeable offset when extrapolated down to zero growth. In the model by Maitra and Dill (2015), this result relies on the turnover of degraded non-ribosomal proteins, whereas in the models by Weiße et al. (2015) and Liao et al. (2017) this behaviour emerges as ribosomes not actively translating mRNAs are more abundant at slow growth.

While these five coarse-grained resource allocation models all provide a better understanding of growth-rate-correlated protein abundance, the role of non-metabolic “housekeeping” proteins is less clear. Maitra and Dill (2015) and Molenaar et al. (2009) disregarded these proteins entirely. Weiße et al. (2015) explicitly included negative autoregulation of housekeeping proteins to approximate constant expression levels across conditions. For Liao et al. (2017), the housekeeping proteins were expressed constitutively and their proteome fraction was only approximately constant. Lastly, Bertaux et al. (2020) fixed the housekeeping allocation manually to a constant fraction.

### 5.2 General considerations for constructing C-GRAMs

Here we provide an overview of the theoretical ideas that underpin our modelling framework. Several excellent reviews of the basic concepts underlying mechanistic whole-cell models have been published in the last few years, notably by De Jong et al. (2017), Bruggeman et al. (2020), and Dourado and Lercher (2020). Such models describe the growing cell as a well-mixed solution of proteins, sometimes transcripts, and intermediate metabolites. The underlying concepts can be used to describe genome-wide models as well as coarse-grained whole-cell models (C-GRAMs) such as ours. What follows here is a synthesis of the concepts as they apply to C-GRAMs.

#### 5.2.1 The dynamics of relative abundances

Consider the vector **X** of absolute abundances associated with all the molecular species present in a given cell. These abundances can be measured in multiple ways; three possible choices are outlined in Table 3. In the C-GRAM presented in this paper, we interpreted the absolute abundances of molecular species i as the mass. Further consider the system size

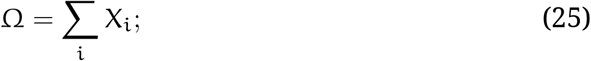

this is equal to the dry mass in our interpretation. For cultures in balanced growth, the growth rate

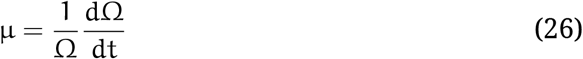

is well-defined and constant. Notably, in this formalism, the growth rate is an emergent property of the system.

The relative abundances (here interpreted as biomass fractions)

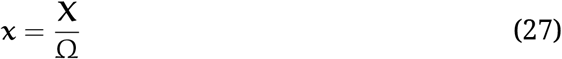

follow the concentration constraint

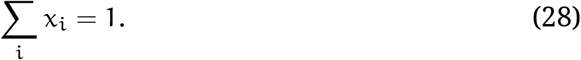

The dynamics of any given cellular constituent i, written in terms of its relative abundance *x*_i_, is calculated as

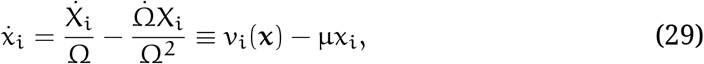

where the net production rate

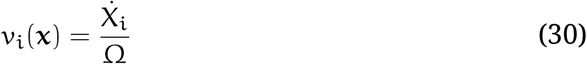

has been written in terms of the relative abundances to create a closed system of equations.

Balanced growth is characterized by the steady state of equation (29, i.e. *ẋ* = 0, such that for each molecular species i, its production rate is balanced by the dilution due to growth:

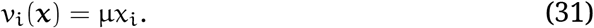

In other words, the relative abundances are constant during balanced growth, because all constituents in the vector **X** grow exponentially with the same rate μ. Furthermore, any well-defined choice of abundance measure must by its construction result in the same growth rate μ.

#### 5.2.2 The interpretation of coarse-grained models

The key choices in formulating a C-GRAM are the biological interpretations of the molecules represented by the relative abundances **x**, together with the functional forms that the fluxes **v**(**x**) assume. In a coarse-grained model, few molecules are included—in other words, the dimensionality of **x** is small, typically between 3 and 10. C-GRAMs such as ours usually focus on protein abundances, with metabolites merely resulting in and possibly regulating protein production. Some models additionally include transcription and translation of mRNA. A very minimal example model was initially proposed by Maitra and Dill (2015) and expanded to include ribosome inactivation (Maitra and Dill 2016). Two examples of models including both transcription and translation are those by Weiße et al. (2015) and Liao et al. (2017). Importantly, molecular species in a coarse-grained resource allocation model do not map one-to-one to molecules that might be found in the cell. Rather, they represent the total abundance of whole classes of proteins.

#### 5.2.3 The stoichiometry matrix

The flux vector **v**(**x**) can be written in terms of the stoichiometry matrix S and a vector *j*(**x**) that contains the conversion rates associated with each reaction (Heinrich and Schuster 1996):

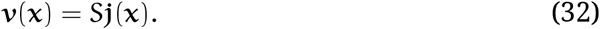

In this matrix S, each column corresponds to a molecular species, and each row to a reaction. Negative values correspond to the consumption of the molecule and positive values correspond to production. If reaction j is balanced, this means that the j-th column of S has sum zero:

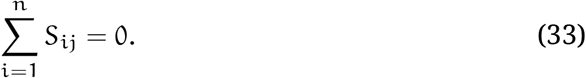

In coarse-grained models, not all reactions are balanced, because the system describes the state of the cell, or of the whole culture. Therefore, reactions exist that grow the cell (or culture) by consuming external nutrients, thereby increasing biomass. For such reactions,

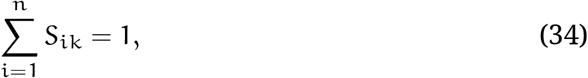

representing that each “firing” of the reaction k increases the total abundance Ω by one unit.

#### 5.2.4 The distinction between proteins and metabolites

At this point, it is helpful to distinguish metabolites from macromolecules. For proteins, v_i_ typically represents only the production rate, because many proteins are stable for many cell generations, and their degradation rates are negligible compared to the dilution rate μ that is due to growth. For metabolites, v_i_ represents the net metabolic flux, incorporating both their production and utilisation. It is usually appropriate to neglect the growth-dilution term for metabolites, because their dynamics are dominated by fast turnover rates. For notation purposes, we split the concentration vector **x** into three parts, representing amino acids, other metabolites, and proteins:

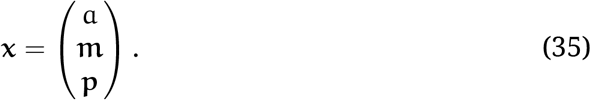

The proteins **p** are the main constituents of the total biomass and are produced from the amino acids a by a single reaction, catalysed by the ribosomes. The remaining metabolites **m** are intermediates. Likewise, the flux vector is decomposed as

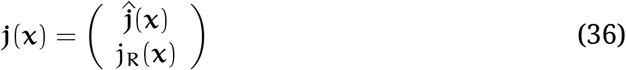

and the flux through the ribosomes j_R_ is separated from the remaining fluxes 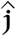.

The protein production reactions are taken to be catalysed by the ribosomes, themselves represented as proteins in this formalism. Importantly, they consume the amino acids a. The relative protein production rates are determined by an allocation vector **f**, which has the same number of components as **p**. Considering a stable protein i that does not change or degrade after its production, its time evolution is given by

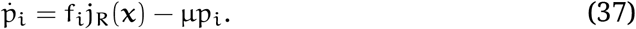

If required, this equation can be supplemented with a degradation term, but in our model we assumed that the proteins were stable. The ribosomal production reactions satisfy mass conservation typified by equation (33). In other words, the consumed substrate a must be balanced with the proteins produced in the ribosomal flux j_R_(**x**). In practice, this means that

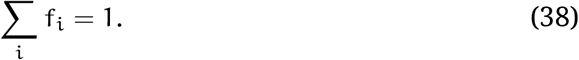

We refer to this as the allocation constraint.

#### 5.2.5 Decomposing the stoichiometry matrix and the dynamic equations

If we assume that all proteins are produced by ribosomes, and do not change afterwards, the stoichiometry matrix S takes a particular form. In a system with n conversion reactions, one amino-acid component, m metabolites, and p proteins, S is an (1 + m + p) by n matrix with the following shape:

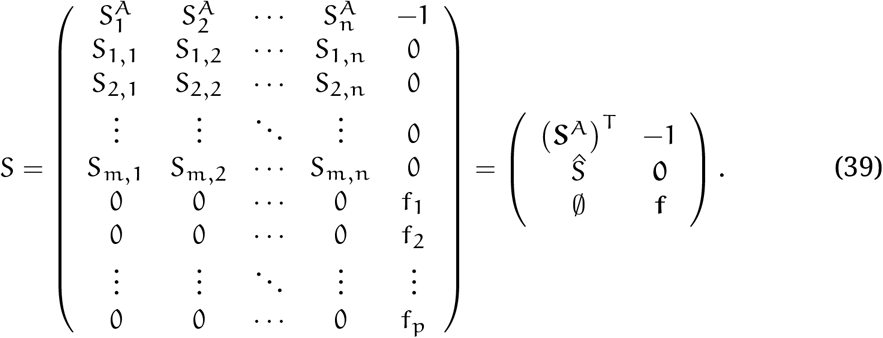

We call the m by n matrix ŝ the metabolic stoichiometry matrix; the (row) vector **S**^*A*^ quantifies the stoichiometry of the reactions that produce the amino acids.

In the end, then, the concentration dynamics of coarse-grained models with a single biomass production reaction catalysed by ribosomes and stable proteins can generally be described as:

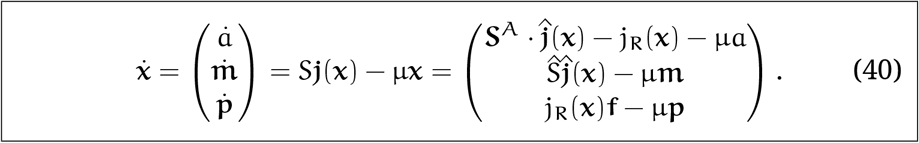

This decomposition is a helpful tool to understand models like the one proposed in this paper, but is is not always applicable. In particular, it does not apply the model by Bertaux et al. (2020), which incorporates conversions between different protein classes after their production.

### 5.3 Enzyme kinetics

The functional forms of the fluxes in the model, as in equation (4), were inspired by models of enzyme kinetics, in particular Michaelis–Menten rate laws. For a Michaelis-Menten enzyme X with concentration *e*_X_, the production rate j^(X)^ of a metabolite by that enzyme through conversion of a substrate with concentration *x*_i_ is given by

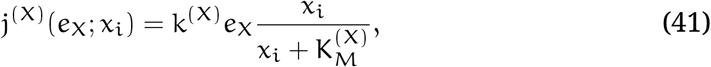

with k^(X)^ and 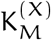 constants that can be related to the enzyme kinetic rates. While there is no direct justification as to why single-enzyme models can be directly applied to the abstracted enzyme classes of coarse-grained models, the Michaelis-Menten rate law has three useful and important properties, namely it is (i) linear in the enzyme concentration, (ii) linear in the substrate concentration at low concentrations, while it (iii) saturates with increasing substrate concentrations.

Empirically, the growth rate of bacterial cultures follows a similar dependency on the availability of a single limiting nutrient source, the concentration of the nutrient source in the environment taking the place of the substrate concentration in the cell (Monod 1949). The dependency from equation (41) is then known as the Monod curve; it can be explained by assuming Michaelis–Menten kinetics of a limiting enzyme (De Jong et al. 2017). However, the empirical observation is strong: the enzyme kinetics described by equation (41) can be used to describe whole pathways, even though pathway fluxes are not always limited by the kinetics of a single enzyme.

For reactions involving more than one substrate, we opted to use a functional form that can be derived using similar assumptions to those underlying Michaelis–Menten kinetics. This meant that the the three useful properties (i)–(iii) noted in the above were maintained. In principle, the flux equation that best describes two-substrate enzyme kinetics depends on the reaction mechanism (Marangoni 2003, ch. 7). A main distinction exists between the ping-pong mechanism, where a product is released before the second substrate binds to the enzyme, and sequential mechanisms, where both substrates bind sequentially, either in a random or an ordered fashion, before any product is released. A general form for the conversion rate j_i_, catalysed by enzyme X with concentration *e*_X_, taking two substrates with concentrations *x*_j_ and x_k_ is given by

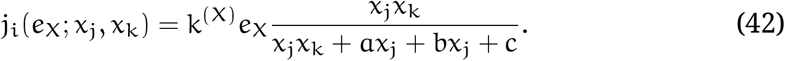

Here k^(X)^ a, b, and c are free constants whose values can be expressed in terms of kinetic rate parameters. Depending on the reaction mechanism, a, b, or c may be equal to zero. It can be seen that the rate function for two-substrate reactions reverts to a Michaelis–Menten dependency when one substrate concentration is held fixed regardless of the reaction mechanism.

In order to compare the parametrizations of one- and two-substrate enzymes, we recast the formula in terms of different parameters as

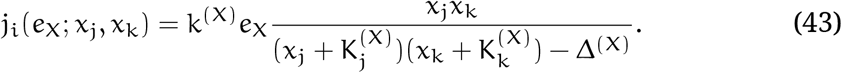

For the simple Δ^(X)^ = 0, the rate law is simply the product of two Michaelis–Menten rate laws. This symmetrical case corresponds to a mechanism where the equilibrium constant for the binding of each of the two substrates does not depend on the presence or absence of the other. In coarse-grained models, reaction mechanisms are not well defined, because the enzymes typically represent entire pathways. We therefore chose Δ__A__f__ = Δ_*A*_r__ = 0 in our model explorations, because this choice treated both substrates symmetrically.

### 5.4 Ribosomal allocation proportional to amino-acid abundance implies the ribosomal growth law

Here, we derive the ribosomal growth law for a class of C-GRAMs that assume that the ribosome allocation depends linearly on the metabolite concentration with an offset equal to the scaled Michaelis constant. This assumption means that

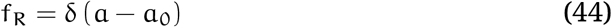

is satisfied dynamically, with δ and a_0_ fixed parameters. In these C-GRAMs, the remaining components of the allocation vector must be chosen such that the allocation constraint

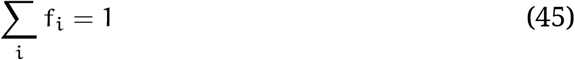

is satisfied. Furthermore, we require that all biomass is produced by stable ribosomes with Michaelis–Menten kinetics, such that the evolution of the ribosomal abundance is given by the ODE

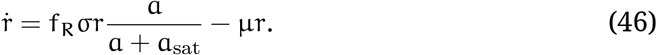

This means that

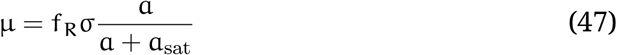

in steady-state (ṙ = 0). Note that the latter two requirements are also met in our original model where ribosomal allocation was growth-optimised alongside the other protein abundances (and, indeed, in the model by Bertaux et al. (2020)). Combining equations (44) and (47) gives

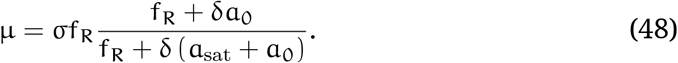

If the ribosomes are saturated, a ≫ a_sat_ and f_R_ is large as well. In this regime, the slope of mu (f_R_) should equal σ according to equation (47). For small values of f_R_, this slope is larger by a factor of 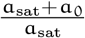. Intuitively, then, fitting a line (with slope σ) to the part of the curve with largef f_R_ will give some nonzero intercept. In the following, the slope and intercept are calculated explicitly.

For a function g(*x*) to approach a straight line given by *C*(*x* – *x*_0_) for large *x*, it means that the following two limits hold:

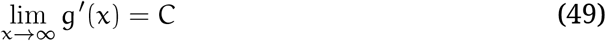

and

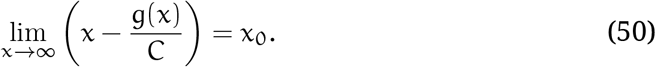

Applying the former to our μ(f_R_) given by equation (48) gives

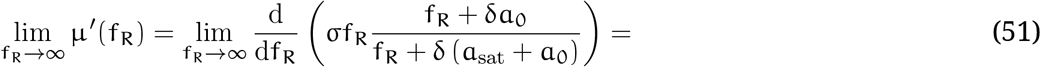

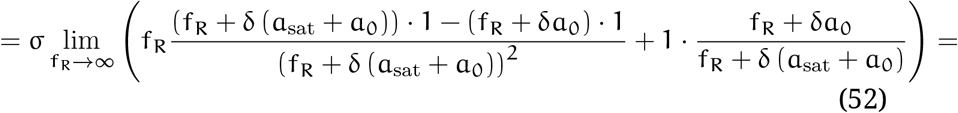

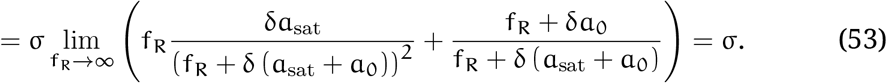

Furthermore,

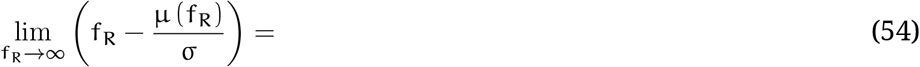

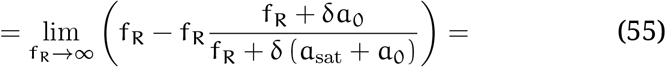

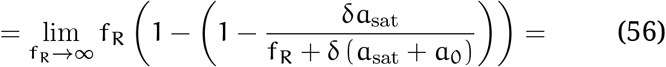

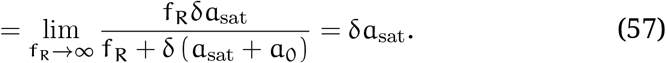

Therefore,

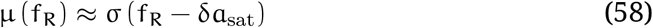

for large f_R_. By inverting this relation and equating the allocation fraction f_R_ to the proteome mass fraction ϕ_R_ in balanced growth, we derive that

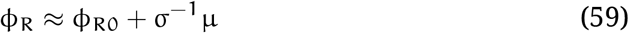

with

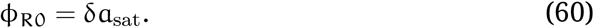

This is the familiar ribosomal growth law.

## 6 Supplementary Tables

## References

Berkhout, Jan, Evert Bosdriesz, Emrah Nikerel, Douwe Molenaar, Dick de Ridder et al. (2013). ‘How Biochemical Constraints of Cellular Growth Shape Evolutionary Adaptations in Metabolism’. In: Genetics 194.2, pp. 505–512. DOI: 10.1534/genetics.113.150631.

Bertaux, François, Julius von Kügelgen, Samuel Marguerat and Vahid Shahrez-aei (2020). ‘A Bacterial Size Law Revealed by a Coarse-Grained Model of Cell Physiology’. In: PLOS Computational Biology 16.9, e1008245. DOI: 10.1371/journal.pcbi.1008245.

Bosdriesz, Evert, Douwe Molenaar, Bas Teusink and Frank J. Bruggeman (2015). ‘How Fast-Growing Bacteria Robustly Tune Their Ribosome Concentration to Approximate Growth-Rate Maximization’. In: FEBS Journal 282.10, pp. 2029–2044. DOI: 10.1111/febs.13258.

Brauer, Matthew J., Curtis Huttenhower, Edoardo M. Airoldi, Rachel Rosenstein, John C. Matese et al. (2008). ‘Coordination of Growth Rate, Cell Cycle, Stress Response, and Metabolic Activity in Yeast’. In: Molecular Biology of the Cell 19.1, pp. 352–367. DOI: 10.1091/mbc.E07-08-0779.

Bremer, Hans and Patrick P. Dennis (2008). ‘Modulation of Chemical Composition and Other Parameters of the Cell at Different Exponential Growth Rates’. In: EcoSal Plus 3.1. DOI: 10.1128/ecosal.5.2.3.

Bruggeman, Frank J, Robert Planqué, Douwe Molenaar and Bas Teusink (2020). ‘Searching for Principles of Microbial Physiology’. In: FEMS Microbiology Reviews 44.6, pp. 821–844. DOI: 10.1093/femsre/fuaa034.

Carlson, C. R., B. Grallert, T. Stokke and E. Boye (1999). ‘Regulation of the Start of DNA Replication in Schizosaccharomyces Pombe’. In: Journal of Cell Science 112.6, pp. 939–946.

Cheng, Chuankai, Edward J. O’Brien, Douglas McCloskey, Jose Utrilla, Connor Olson et al. (2019). ‘Laboratory Evolution Reveals a Two-Dimensional Rate-Yield Tradeoff in Microbial Metabolism’. In: PLOS Computational Biology 15.6, e1007066. DOI: 10.1371/journal.pcbi.1007066.

Christodoulou, Dimitris, Hannes Link, Tobias Fuhrer, Karl Kochanowski, Luca Ge-rosa et al. (2018). ‘Reserve Flux Capacity in the Pentose Phosphate Pathway Enables Escherichia Coli’s Rapid Response to Oxidative Stress’. In: Cell Systems 6.5, 569–578.e7. DOI: 10.1016/j.cels.2018.04.009.

Chubukov, Victor, Luca Gerosa, Karl Kochanowski and Uwe Sauer (2014). ‘Co-ordination of Microbial Metabolism’. In: Nature Reviews Microbiology 12.5, pp. 327–340. DOI: 10.1038/nrmicro3238.

Dai, Xiongfeng and Manlu Zhu (2020). ‘Coupling of Ribosome Synthesis and Translational Capacity with Cell Growth’. In: Trends in Biochemical Sciences 45.8, pp. 681–692. DOI: 10.1016/j.tibs.2020.04.010.

Dai, Xiongfeng, Manlu Zhu, Mya Warren, Rohan Balakrishnan, Vadim Patsalo et al. (2016). ‘Reduction of Translating Ribosomes Enables *Escherichia Coli* to Maintain Elongation Rates during Slow Growth’. In: Nature Microbiology 2.2, p. 2016231. DOI: 10.1038/nmicrobiol.2016.231.

De Jong, Hidde, Stefano Casagranda, Nils Giordano, Eugenio Cinquemani, Delphine Ropers et al. (2017). ‘Mathematical Modelling of Microbes: Metabolism, Gene Expression and Growth’. In: Journal of The Royal Society Interface 14.136, p. 20170502. DOI: 10.1098/rsif.2017.0502.

Dekel, Erez and Uri Alon (2005). ‘Optimality and Evolutionary Tuning of the Expression Level of a Protein’. In: Nature 436.7050, nature03842. DOI: 10.1038/nature03842.

De Queiroz, José Humberto, Jean-Louis Uribelarrea and Alain Pareilleux (1993). ‘Estimation of the Energetic Biomass Yield and Efficiency of Oxidative Phosphorylation in Cell-Recycle Cultures of Schizosaccharomyces Pombe’. In: Applied Microbiology and Biotechnology 39.4, pp. 609–614. DOI: 10.1007/BF00205061.

Dourado, Hugo and Martin J. Lercher (2020). ‘An Analytical Theory of Balanced Cellular Growth’. In: Nature Communications 11.1 (1), pp. 1–14. DOI: 10.1038/s41467-020-14751-w.

Elsemman, Ibrahim E., Angelica Rodriguez Prado, Pranas Grigaitis, Manuel Garcia Albornoz, Victoria Harman et al. (2022). ‘Whole-Cell Modeling in Yeast Predicts Compartment-Specific Proteome Constraints That Drive Metabolic Strategies’. In: Nature Communications 13.1 (1), p. 801. DOI: 10.1038/s41467-022-28467-6.

Ertugay, N. and H. Hamamci (1997). ‘Continuous Cultivation of Bakers’ Yeast: Change in Cell Composition at Different Dilution Rates and Effect of Heat Stress on Trehalose Level’. In: Folia Microbiologica 42.5, pp. 463–467. DOI: 10.1007/BF02826554.

Fantes, P. and P. Nurse (1977). ‘Control of Cell Size at Division in Fission Yeast by a Growth-Modulated Size Control over Nuclear Division’. In: Experimental Cell Research 107.2, pp. 377–386. DOI: 10.1016/0014-4827(77)90359-7.

Gao, Fuchang and Lixing Han (2012). ‘Implementing the Nelder-Mead Simplex Algorithm with Adaptive Parameters’. In: Computational Optimization and Applications 51.1, pp. 259–277. DOI: 10.1007/s10589-010-9329-3.

Godard, Patrice, Antonio Urrestarazu, Stéphan Vissers, Kevin Kontos, Gianluca Bontempi et al. (2007). ‘Effect of 21 Different Nitrogen Sources on Global Gene Expression in the Yeast Saccharomyces Cerevisiae’. In: Molecular and Cellular Biology 27.8, pp. 3065–3086. DOI: 10.1128/MCB.01084-06.

Grigaitis, Pranas, Douwe A. J. Grundel, Eunice van Pelt-KleinJan, Mirushe Isaku, Guixiang Xie et al. (2022). ‘A Computational Toolbox to Investigate the Metabolic Potential and Resource Allocation in Fission Yeast’. In: mSystems. DOI: 10.1128/msystems.00423-22.

Harris, Leigh K. and Julie A. Theriot (2016). ‘Relative Rates of Surface and Volume Synthesis Set Bacterial Cell Size’. In: Cell 165.6, pp. 1479–1492. DOI: 10.1016/j.cell.2016.05.045.

Heinrich, Reinhart and Stefan Schuster (1996). The Regulation of Cellular Systems. New York, NY: Chapman & Hall. 372 pp.

Hu, Xiao-Pan, Hugo Dourado, Peter Schubert and Martin J. Lercher (2020). ‘The Protein Translation Machinery Is Expressed for Maximal Efficiency in Escherichia Coli’. In: Nature Communications 11.1 (1), p. 5260. DOI: 10.1038/s41467-020-18948-x.

Hui, Sheng, Josh M. Silverman, Stephen S. Chen, David W. Erickson, Markus Basan et al. (2015). ‘Quantitative Proteomic Analysis Reveals a Simple Strategy of Global Resource Allocation in Bacteria’. In: Molecular Systems Biology 11, p. 784.

Hurbain, Julien, Quentin Thommen, Francois Anquez and Benjamin Pfeuty (2022). ‘Quantitative Modeling of Pentose Phosphate Pathway Response to Oxidative Stress Reveals a Cooperative Regulatory Strategy’. In: iScience 25.8, p. 104681. DOI: 10.1016/j.isci.2022.104681.

Jacoby, Richard P., Antonella Succurro and Stanislav Kopriva (2020). ‘Nitrogen Substrate Utilization in Three Rhizosphere Bacterial Strains Investigated Using Proteomics’. In: Frontiers in Microbiology 11.

Kleijn, Istvan T., Amalia Martínez-Segura, François Bertaux, Malika Saint, Holger Kramer et al. (2022). ‘Growth-Rate-Dependent and Nutrient-Specific Gene Expression Resource Allocation in Fission Yeast’. In: Life Science Alliance 5.5, e202101223. DOI: 10.26508/lsa.202101223.

Koch, Arthur L. (1988). ‘Why Can’t a Cell Grow Infinitely Fast?’ In: Canadian Journal of Microbiology 34.4, pp. 421–426. DOI: 10.1139/m88-074.

Liao, Chen, Andrew E. Blanchard and Ting Lu (2017). ‘An Integrative Circuit–Host Modelling Framework for Predicting Synthetic Gene Network Behaviours’. In: Nature Microbiology 2, pp. 1658–1666. DOI: 10.1038/s41564-017-0022-5.

López-Maury, Luis, Samuel Marguerat and Jürg Bähler (2008). ‘Tuning Gene Expression to Changing Environments: From Rapid Responses to Evolutionary Adaptation’. In: Nature Reviews Genetics 9.8, pp. 583–593. DOI: 10.1038/nrg2398.

Lynch, Michael, Mark C. Field, Holly V. Goodson, Harmit S. Malik, José B. Pereira-Leal et al. (2014). ‘Evolutionary Cell Biology: Two Origins, One Objective’. In: Proceedings of the National Academy of Sciences 111.48, pp. 16990–16994. DOI: 10.1073/pnas.1415861111.

Maitra, Arijit and Ken A. Dill (2015). ‘Bacterial Growth Laws Reflect the Evolutionary Importance of Energy Efficiency’. In: Proceedings of the National Academy of Sciences 112.2, pp. 406–411. DOI: 10.1073/pnas.1421138111.

Maitra, Arijit and Ken A. Dill (2016). ‘Modeling the Overproduction of Ribosomes When Antibacterial Drugs Act on Cells’. In: Biophysical Journal 110.3, pp. 743–748. DOI: 10.1016/j.bpj.2015.12.016.

Malecki, Michal, Stephan Kamrad, Markus Ralser and Jürg Bähler (2020). ‘Mitochondrial Respiration Is Required to Provide Amino Acids during Fermentative Proliferation of Fission Yeast’. In: EMBO reports 21.11, e50845. DOI: 10.15252/embr.202050845.

Marangoni, Alejandro G. (2003). Enzyme Kinetics: A Modern Approach. Hoboken, NJ: John Wiley & Sons, Inc. ISBN: 978-0-471-15985-8.

Metzl-Raz, Eyal, Moshe Kafri, Gilad Yaakov, Ilya Soifer, Yonat Gurvich et al. (2017). ‘Principles of Cellular Resource Allocation Revealed by Condition-Dependent Proteome Profiling’. In: eLife 6, e28034. DOI: 10.7554/eLife.28034.

Milo, Ron, Paul Jorgensen, Uri Moran, Griffin Weber and Michael Springer (2010). ‘BioNumbers—the Database of Key Numbers in Molecular and Cell Biology’. In: Nucleic Acids Research 38 (suppl_1), pp. D750–D753. DOI: 10.1093/nar/gkp889.

Mitchison, J. M. and K. G. Lark (1962). ‘Incorporation of 3H-adenine into RNA during the Cell Cycle of Schizosaccharomyces Pombe’. In: Experimental Cell Research 28.2, pp. 452–455. DOI: 10.1016/0014-4827(62)90304-X.

Mogensen, Patrick K. and Asbjørn N. Riseth (2018). ‘Optim: A Mathematical Optimization Package for Julia’. In: Journal of Open Source Software 3.24, p. 615. DOI: 10.21105/joss.00615.

Molenaar, Douwe, Rogier van Berlo, Dick de Ridder and Bas Teusink (2009). ‘Shifts in Growth Strategies Reflect Tradeoffs in Cellular Economics’. In: Molecular Systems Biology 5.1, p. 323. DOI: 10.1038/msb.2009.82.

Monod, Jacques (1949). ‘The Growth of Bacterial Cultures’. In: Annual Review of Microbiology 3.1, pp. 371–394. DOI: 10.1146/annurev.mi.03.100149.002103.

Mori, Matteo, Severin Schink, David W. Erickson, Ulrich Gerland and Terence Hwa (2017). ‘Quantifying the Benefit of a Proteome Reserve in Fluctuating Environments’. In: Nature Communications 8.1, p. 1225. DOI: 10.1038/s41467-017-01242-8.

Neidhardt, Frederick C., John L. Ingraham and Moselio Schaechter (1990). Physiology of the Bacterial Cell. Sunderland, MA: Sinauer Associates.

Nelder, J. A. and R. Mead (1965). ‘A Simplex Method for Function Minimization’. In: The Computer Journal 7.4, pp. 308–313. DOI: 10.1093/comjnl/7.4.308.

O’Brien, Edward J., Jose Utrilla and Bernhard O. Palsson (2016). ‘Quantification and Classification of E. Coli Proteome Utilization and Unused Protein Costs across Environments’. In: PLOS Computational Biology 12.6, e1004998. DOI: 10.1371/journal.pcbi.1004998.

Odermatt, Pascal D, Teemu P Miettinen, Joël Lemière, Joon Ho Kang, Emrah Bostan et al. (2021). ‘Variations of Intracellular Density during the Cell Cycle Arise from Tip-Growth Regulation in Fission Yeast’. In: eLife 10, e64901. DOI: 10.7554/eLife.64901.

Ojkic, Nikola, Diana Serbanescu and Shiladitya Banerjee (2019). ‘Surface-to-Volume Scaling and Aspect Ratio Preservation in Rod-Shaped Bacteria’. In: eLife 8, e47033. DOI: 10.7554/eLife.47033.

Oldewurtel, Enno R., Yuki Kitahara, Baptiste Cordier, Gizem Özbaykal and Sven van Teeffelen (2019). ‘Bacteria Control Cell Volume by Coupling Cell-Surface Expansion to Dry-Mass Growth’. In: bioRxiv, p. 769786. DOI: 10.1101/769786.

Pandey, Parth Pratim and Sanjay Jain (2016). ‘Analytic Derivation of Bacterial Growth Laws from a Simple Model of Intracellular Chemical Dynamics’. In: Theory in Biosciences 135.3, pp. 121–130. DOI: 10.1007/s12064-016-0227-9.

Peebo, Karl, Kaspar Valgepea, Andres Maser, Ranno Nahku, Kaarel Adamberg et al. (2015). ‘Proteome Reallocation in Escherichia Coli with Increasing Specific Growth Rate’. In: Molecular BioSystems 11.4, pp. 1184–1193. DOI: 10.1039/C4MB00721B.

Planqué, Robert, Josephus Hulshof, Bas Teusink, Johannes C. Hendriks and Frank J. Bruggeman (2018). ‘Maintaining Maximal Metabolic Flux by Gene Expression Control’. In: PLOS Computational Biology 14.9, e1006412. DOI: 10.1371/journal.pcbi.1006412.

Rackauckas, Christopher and Qing Nie (2017). ‘DifferentialEquations.Jl – A Performant and Feature-Rich Ecosystem for Solving Differential Equations in Julia’. In: Journal of Open Research Software 5.1. DOI: 10.5334/jors.151.

Schaechter, M., O. Maaløe and N. O. Kjeldgaard (1958). ‘Dependency on Medium and Temperature of Cell Size and Chemical Composition during Balanced Growth of Salmonella Typhimurium’. In: Journal of General Microbiology 19.3, pp. 592–606. DOI: 10.1099/00221287-19-3-592.

Schaechter, Moselio (2006). ‘From Growth Physiology to Systems Biology’. In: International Microbiology: The Official Journal of the Spanish Society for Microbiology 9.3, pp. 157–161.

Schmidt, Alexander, Karl Kochanowski, Silke Vedelaar, Erik Ahrné, Benjamin Volkmer et al. (2016). ‘The Quantitative and Condition-Dependent Escherichia Coli Proteome’. In: Nature Biotechnology 34.1, pp. 104–110. DOI: 10.1038/nbt.3418.

Scott, Matthew, Carl W. Gunderson, Eduard M. Mateescu, Zhongge Zhang and Terence Hwa (2010). ‘Interdependence of Cell Growth and Gene Expression: Origins and Consequences’. In: Science 330, pp. 1099–1102. DOI: 10.1126/science.1192588.

Scott, Matthew, Stefan Klumpp, Eduard M. Mateescu and Terence Hwa (2014). ‘Emergence of Robust Growth Laws from Optimal Regulation of Ribosome Synthesis’. In: Molecular Systems Biology 10.8, p. 747. DOI: 10.15252/msb.20145379.

Towbin, Benjamin D., Yael Korem, Anat Bren, Shany Doron, Rotem Sorek et al. (2017). ‘Optimality and Sub-Optimality in a Bacterial Growth Law’. In: Nature Communications 8, p. 14123. DOI: 10.1038/ncomms14123.

Valgepea, Kaspar, Kaarel Adamberg, Andrus Seiman and Raivo Vilu (2013). ‘Escherichia Coli Achieves Faster Growth by Increasing Catalytic and Translation Rates of Proteins’. In: Molecular BioSystems 9.9, pp. 2344–2358. DOI: 10.1039/C3MB70119K.

Von Stockar, U. and J. -S. Liu (1999). ‘Does Microbial Life Always Feed on Negative Entropy? Thermodynamic Analysis of Microbial Growth’. In: Biochimica et Bio-physica Acta (BBA) - Bioenergetics 1412.3, pp. 191–211. DOI: 10.1016/S0005-2728(99)00065-1.

Waldron, C. and F. Lacroute (1975). ‘Effect of Growth Rate on the Amounts of Ribosomal and Transfer Ribonucleic Acids in Yeast.’ In: Journal of Bacteriology 122.3, pp. 855–865.

Weiße, Andrea Y., Diego A. Oyarzún, Vincent Danos and Peter S. Swain (2015). ‘Mechanistic Links between Cellular Trade-Offs, Gene Expression, and Growth’. In: Proceedings of the National Academy of Sciences 112.9, E1038–E1047. DOI: 10.1073/pnas.1416533112.

Wu, Chenhao, Rohan Balakrishnan, Nathan Braniff, Matteo Mori, Gabriel Man-zanarez et al. (2022). ‘Cellular Perception of Growth Rate and the Mechanistic Origin of Bacterial Growth Law’. In: Proceedings of the National Academy of Sciences of the United States of America 119.20, e2201585119. DOI: 10.1073/pnas.2201585119.

You, Conghui, Hiroyuki Okano, Sheng Hui, Zhongge Zhang, Minsu Kim et al. (2013). ‘Coordination of Bacterial Proteome with Metabolism by Cyclic AMP Signalling.’ In: Nature 500, pp. 301–306. DOI: 10.1038/nature12446.

Yu, Rosemary, Kate Campbell, Rui Pereira, Johan Björkeroth, Qi Qi et al. (2020). ‘Nitrogen Limitation Reveals Large Reserves in Metabolic and Translational Capacities of Yeast’. In: Nature Communications 11.1 (1), pp. 1–12. DOI: 10.1038/s41467-020-15749-0.

Yu, Rosemary, Egor Vorontsov, Carina Sihlbom and Jens Nielsen (2021). ‘Quantifying Absolute Gene Expression Profiles Reveals Distinct Regulation of Central Carbon Metabolism Genes in Yeast’. In: eLife 10, e65722. DOI: 10.7554/eLife.65722.

Zavřel, Tomáš, Marjan Faizi, Cristina Loureiro, Gereon Poschmann, Kai Stühler et al. (2019). ‘Quantitative Insights into the Cyanobacterial Cell Economy’. In: eLife 8, e42508. DOI: 10.7554/eLife.42508.

